# Gene expression and eQTL analysis reflect the heterogeneity in the inflammatory status of the duodenal epithelial lining in coeliac disease

**DOI:** 10.1101/2024.02.29.582756

**Authors:** Aarón D. Ramírez-Sánchez, Stephanie Zühlke, Raúl Aguirre-Gamboa, Martijn Vochteloo, Lude Franke, Knut E.A. Lundin, Sebo Withoff, Iris H. Jonkers

## Abstract

In coeliac disease (CeD), the epithelial lining (EL) of the small intestine is severely damaged through a complex auto-inflammatory response that results in intraepithelial lymphocytes (IELs) attacking epithelial cells (ECs). To better understand the changes occurring in the EL in CeD, including after initiation of a gluten-free diet, and the regulatory mechanisms that affect mucosal homeostasis, we investigated ECs and IELs in the CeD duodenal EL using RNA-seq and eQTL analysis on predicted cell types. The study included 82 duodenal biopsies from volunteers, grouped into controls (CTRL), gluten-free diet treated CeD (TCD) and untreated CeD (UCD). We identified 2,868 differential expressed genes, which clustered into four sets with specific functions related to CeD. Two sets, one upregulated for cell cycle function and one downregulated for digestion, transmembrane transport, and laminin pathways, defined three groups of samples based on inflammation status: non-inflamed, mild inflammation or severe inflammation. These inflammation states correlated with but were not specific to disease state. The remaining two sets of genes were enriched for immune, extracellular matrix, and barrier functions. These latter two sets allowed the classification of samples into their disease conditions: CTRL, TCD, and UCD. Finally, deconvoluting eQTL effects from ECs and immune cells in the EL identified 7 and 9 cell-type-mediated eQTL genes, respectively. In sum, we identified genes expressed in the duodenal EL whose expression might be used as biomarkers to assess CeD condition and its mucosal and immune status.

**Highlights:**

- Gene expression in epithelial lining in CeD patients reflects inflammation status
- The transcriptome can classify epithelium as no, mild, or severe inflammation
- Cell cycle, absorption, and basal lamina genes are indicators of mucosal damage
- Immune and extracellular matrix genes distinguish untreated from treated cases
- Predicted-cell-type eQTLs pinpoint effects in immune and epithelial cells

## Introduction

Celiac Disease (CeD) is a complex immune-mediated disorder caused by the intake of dietary gluten, a protein found in wheat, barley, and rye, in individuals with a genetic predisposition [1]. The main tissue affected in CeD is the epithelium of the duodenum, which consists of two main compartments – the epithelial lining (EL) and the lamina propria – separated by the basal lamina. The EL is a single cell layer characterised by finger-like structures called villi and invaginations called crypts. It comprises multiple epithelial cell (EC) types, including enterocytes, Tuft cells, goblet cells, enteroendocrine cells, crypt-residing adult stem cells and Paneth cells, which together account for ∼90% of cells in the EL [2]. In CeD, the villus structure of the epithelium is affected and characterised by villus atrophy and crypt hyperplasia [3]. Moreover, in CeD the EL is invaded by immune cells designated intraepithelial lymphocytes (IELs) [4].

In CeD, IELs gain a cytotoxic phenotype resulting from a complex immune reaction. First, dietary gluten is partially digested into gliadin peptides, which are deamidated by tissue transglutaminase 2 [5–8]. In the lamina propria, these deamidated gliadin peptides are then presented by HLA-DQ2-and/or -DQ8-expressing antigen-presenting cells that are recognised by gluten-specific CD4+ T cells, causing the latter population to expand [9]. The gluten-specific CD4+ T cells in turn promote the development of B cells and the production of antibodies, and they activate CD8+ T cells that move to the EL and develop into IELs [4,10]. IEL populations are categorised into CD4 or CD8 subclasses based on the composition of their T cell receptor (TCR). The IEL populations present in the gut are TCRαβ+CD8αβ+, TCRαβ+CD4+, and TCRγδ+ T cells. In CeD, TCRαβ+CD8αβ+ IELs acquire a lymphokine killer-like activity by aberrantly expressing NK-lineage genes, including killer cell lectin-like receptor C2 (*KLRC2*, also known as *NKG2C*), natural cytotoxicity triggering receptor 1 (*NCR1*, also known as *NKp46*), and *NCR2* (also known as *NKp44*). The current thinking is that these cytotoxic TCRαβ+CD8αβ+ IELs cause the EC damage in CeD [4,11–17].

Once CeD patients start a gluten-free diet (GFD), some symptoms may alleviate within weeks, but overall mucosal recovery varies between patients and is only achieved in half of CeD patients after one year of GFD [18]. Crypt hyperplasia and villus atrophy gradually recover over time after gluten is excluded from the diet, and immune cells implicated in CeD pathogenesis, like gluten-specific TCRαβ+ CD4^+^ T cells in the lamina propria and cytolytic TCRαβ+ CD8+ IELs, decrease in numbers [19]. Remarkably, however, TCRγδ+ IELs remain elevated even after inflammation is dampened [20,21]. Inflammation associated to CeD also permanently reconfigures TCRγδ+ IELs, shifting them from a cell population that maintains homeostasis in the local environment of the gut to an altered cell population producer of IFNy [22]. Understanding the causes of variation in CeD severity and recovery can therefore improve our ability to identify the underlying pathways that lead to disease (and repair) and help identify biomarkers suggestive of active disease and mucosal recovery.

To better understand the changes occurring in the EL in CeD, including after GFD, and the regulatory mechanisms that affect mucosal homeostasis in CeD, we investigated ECs and IELs in the CeD duodenal EL using RNA-seq and predicted cell-type eQTL analysis. Using gene expression profiles of samples, we could distinguish three EL inflammation states: non-inflamed, mild inflammation, and severe inflammation, and these inflammation states correlated with but were not specific to disease state. We further analysed gene expression to determine their gene function in CeD and their interaction with SNPs associated to CeD.

## Results

### Marsh score and disease condition are the main drivers of the transcriptomic landscape

To study the transcriptional heterogeneity in CeD, we analysed EL isolated from intestinal biopsies from 90 adult individuals, classified into three groups based on their condition: controls (CTRL, volunteers undergoing upper endoscopy for complaints unrelated to CeD; n=30), treated CeD cases (TCD, CeD patients under GFD with a Marsh score ≤ 2; n=30) and untreated CeD cases (UCD, CeD patients previously diagnosed or suspected of having CeD with a Marsh score of 3 and/or clinical manifestations of CeD; n=30). From these biopsy samples, we isolated cells for subsequent flow cytometry analysis and RNA extraction. Extracted RNA was used to generate poly(A)-RNA-seq libraries, followed by comprehensive sequencing. Eight outliers were excluded from the dataset as they met at least three of the following criteria: outliers in principal component analysis (PCA), unique pair read percentages < 10%, unmapped read percentages > 10%, RNA concentration < 1.5 ng/µL, total number of reads > 3 million, sample present in batch 9 of RNA extraction, or degradation ratio > 10% (**Supplementary Table 1**). Our analysis then continued with a refined dataset of 25 CTRLs, 28 TCDs, and 29 UCDs (**Table 1**), wherein 20,498 genes were consistently detected across all libraries (**Fig. 1A**).

**Figure 1.**
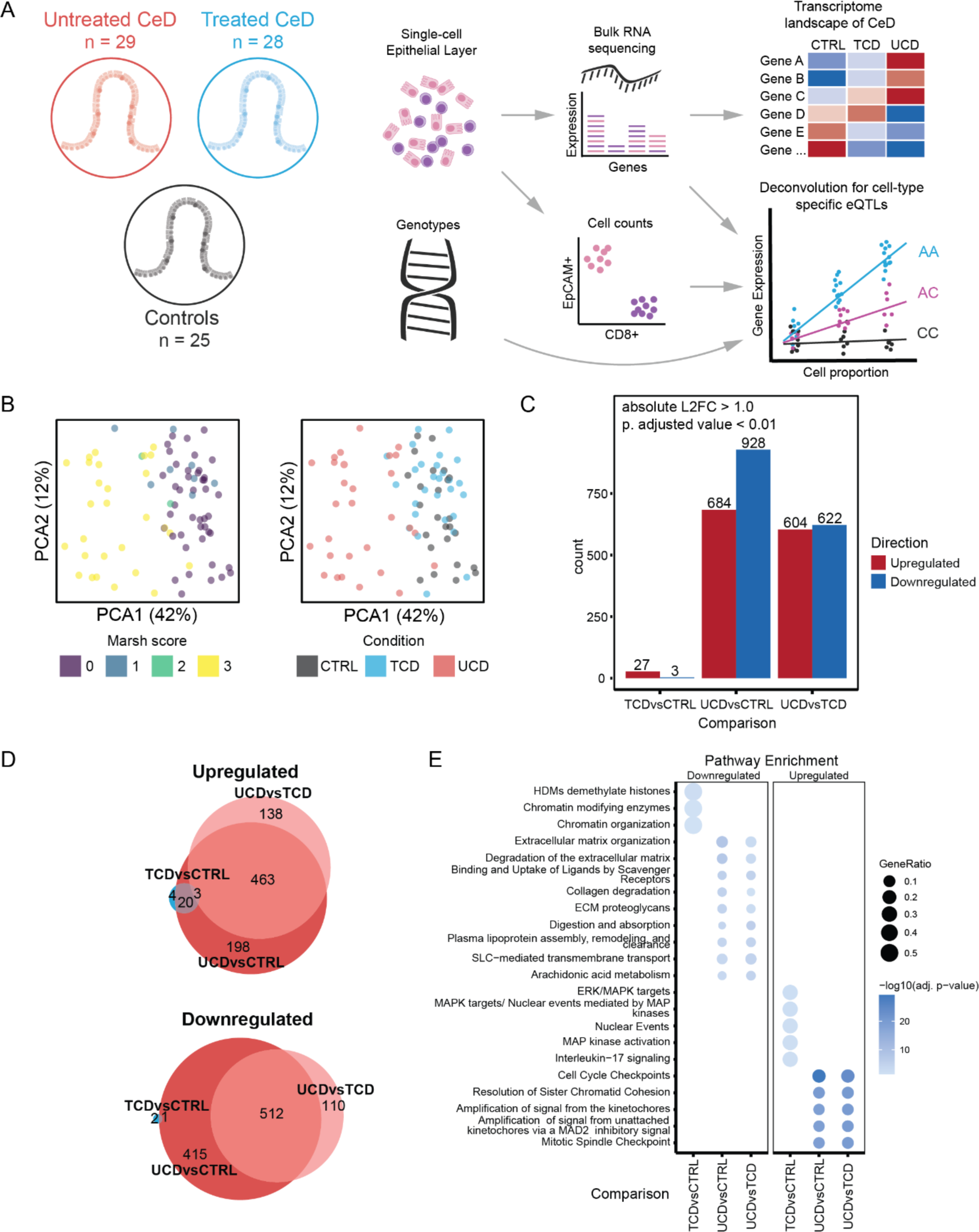
Study design and transcriptome features of the duodenal epithelial lining (EL) in CeD. **(A)** Study design. 82 duodenal biopsies were obtained from volunteers grouped into controls (CTRL), GFD-treated CeD cases (TCD), and untreated CeD cases (UCD). Duodenal biopsies were processed to obtain EL cells in a single-cell suspension. Resulting EL cells were divided to perform cytometry analysis to obtain cell-type counts and to extract bulk-RNA. Genotypes were obtained from whole blood. RNA was processed in poly(A) RNA libraries and paired-end sequenced. Bulk-RNA reads were used to study the transcriptome landscape of CeD by comparing groups. Bulk-RNA expression, cell counts, and genotypes were used to obtain both bulk eQTL and predicted-cell-type eQTL effects. **(B)** Principal component analysis (PCA) of all duodenal samples using the top 1000 most variably expressed genes coloured by Marsh score (left) and CeD condition (right). Each dot corresponds to a single sample. Only the first two principal components are shown. **(C)** Overview of differentially expressed (DE) genes obtained by comparing TCD vs CTRL, UCD vs CTRL, and UCD vs TCD. DE genes are described as upregulated (red) or downregulated (blue). Absolute Log2 Fold change (L2FC) > 1. Adjusted p-value < 0.01. See Supplementary *Table 2* for complete gene lists. **(D)** Overlap of DE genes represented as an Euler diagram. **(E)** The top five most significantly enriched pathways uncovered by pathway enrichment analysis of the DE genes shown in (D) identified using the Reactome database. Dot size indicates the ratio of the number of genes present in the gene set to the total gene set used in each pathway. Dot shading indicates the adjusted p-value after negative log10 transformation.

**Table 1.** Descriptive characteristics of the cohort displayed as number (percentage) or median (interquartile range).See Supplementary table 1 for additional information.

|  | Total cohort | Controls | Treated CeD | Untreated CeD |
| --- | --- | --- | --- | --- |
| **Size** | 82 | 25 | 28 | 29 |
| **Marsh classification** |  |  |  |  |
| **0** | 44 (54%) | 25 (100%) | 19 (68%) |  |
| **1** | 7 (9%) |  | 7 (25%) | 1 (3%) |
| **2** | 2 (2%) |  | 2 (7%) |  |
| **3** | 29 (35%) |  |  | 28 (97%) |
| **Sex** |  |  |  |  |
| **Male** | 36 (44%) | 17 (68%) | 8 (29%) | 11 (38%) |
| **Female** | 46 (56%) | 8 (32%) | 20 (71%) | 18 (62%) |
| **Age (years)** | 44 (32–58) | 43 (31–60) | 45 (36–55) | 41.5 (26–58) |

We performed PCA to assess the variables that explain the transcriptomic landscape. We explored the first 10 principal components (PCs) by correlating them with different variables (**Suppl. Fig. 1-2**). We observed that PC1 was explained by Marsh score, disease classification, and crypt ratio length (calculated as the ratio of expression of apolipoprotein (*APO*) A4 and marker of proliferation (*M*) KI67) [23–25] (**Fig. 1B**). Sex, age, and technical characteristics (sequencing depth, sequencing batch, %GC content, and RNA integrity) were the main determinants for PCs 2–10, and these characteristics were subsequently used as covariates in all analyses (**Suppl. Fig. 1**).

**Figure 2.**
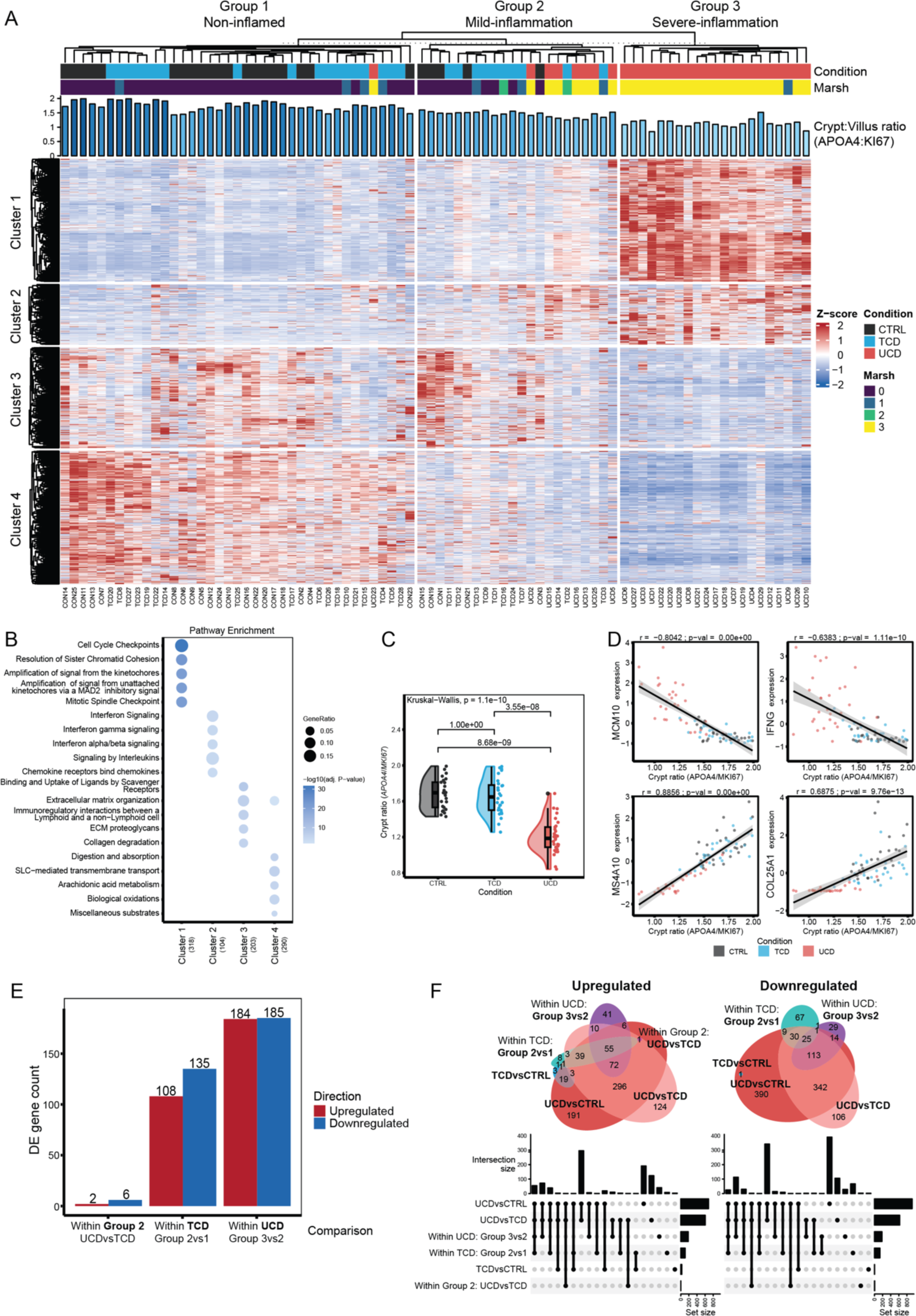
Heterogeneity of the EL transcriptome in CeD allows classification of duodenal biopsies into non-inflamed, mild inflammation, and severe inflammation. (A) Heatmap of gene expression of DE genes in all samples. Each row corresponds to a DE gene. Each column corresponds to a single sample. DE genes clustered into four clusters and three groups. Shown are the CeD condition, Marsh score, and the APOA4:KI67 ratios. Gene expression, scaled and centred around 0, is shown as a Z-score. (B) Pathway enrichment analysis of each cluster of genes (x-axis) using the Reactome database. The five most significant pathways per cluster are shown (y-axis). The number in brackets indicates the number of DE genes present in all enrichments. (C) Statistical comparison of the APOA4:MKI67 ratio, which is an indicator of crypt length (adjusted p-value < 0.01, Dunn test, Bonferroni correction). (D) Correlation plots of DE genes representative of each cluster with APOA4:KI67 ratios. From left to right and top to bottom, each gene is a representative example from cluster 1 to 4, respectively. Each plot shows the correlation between called and centred gene expression and the APOA4:KI67 ratio. Each dot corresponds to a single sample. Dot colour indicates CeD condition. (E) Overview of significant DE genes resulting from comparisons between Group 2 UCD and CTRL, within TCD (Group 2 vs 1), and within UCD (Group 3 vs 2) on the x-axis. Directions of DE are indicated by colour: upregulated (red) and downregulated (blue). See Supplementary Table 4 for complete gene lists. (F) Overlap of DE genes between all tested comparisons. Genes are divided by upregulated (right) and downregulated (left) sets, represented using Euler diagrams (top) and Upset plots (bottom).

To evaluate the effects of CeD and treatment with GFD on gene expression in the EL of the small intestine, we performed differential expression analysis contrasting UCD vs CTRL, TCD vs CTRL, and UCD vs TCD. In total, we identified 2,868 differentially expressed (DE) genes (**Fig. 1C**, absolute Log2 Fold change (L2FC) > 1, adjusted p-value < 0.01) (**Supplementary Table 2**). The UCD vs CTRL comparison showed the highest number of DE genes (n=1,612), followed by UCD vs TCD (n=1,226), whereas TCD vs CTRL exhibited only 30 DE genes. Most of the UCD vs CTRL and UCD vs TCD DE genes (%) overlap and are concordant in direction (**Fig. 1D**, **Suppl. Fig. 3**). Thus, treated CeD and control individuals are similar, whereas the untreated CeD condition is associated with large changes in gene expression in cells present in the EL.

Taking direction into consideration, we performed enrichment analysis to explore the function of the DE genes (**Fig. 1E**, adjusted p-value < 0.05) (**Supplementary Table 3**). As expected, UCD vs CTRL and UCD vs TCD exhibited remarkably similar enriched pathways. Upregulated genes such as cell-division cycle 45 (*CDC45*), minichromosome maintenance 2 (*MCM2*), and origin recognition complex 1 (*ORC1*) caused enrichment for ‘cell cycle pathways’. In the same comparisons, the downregulated genes collagen 4A1 (e.g. *COL4A1*), vitronectin (*VTN*), and laminin A5 (*LAMA5*) caused enrichment for ‘extracellular matrix function’. In addition, the ‘digestion and absorption pathways’ were enriched via downregulation of genes encoding the digestive enzymes lactase (*LCT*) and trehalase (*TREH*), as well as solute carrier family genes (e.g. *SLC2A5*). It is likely that these observations are associated with increased proliferation of IELs and ECs in crypts and loss of differentiated absorptive ECs in the villi, which are both hallmarks of CeD [1,26,27].

For the TCD vs CTRL comparison, we found an enrichment for Mitogen-Activated Protein Kinase (MAPK) pathways based on the upregulated genes (e.g. *MAPK11*). This suggests that, although treated CeD samples resemble control samples, recovery may not be complete, or that TCD samples display persistent alterations in the epithelium because they previously went through an auto-inflammatory state.

### Classification of CeD states into conditions characterised by severely inflamed, mildly inflamed, or recovered epithelium

K-means clustering analysis indicated the presence of three groups in our data set using an optimal k = 3 determined using three different approaches (**Fig. 2A**, **Suppl. Fig. 4**). For this, we used all DE genes and expected to identify groups primarily consisting of CTRLs, TCDs, and UCDs. However, when taking the distribution of samples in each group and the pathway analysis into consideration, we think that the groups can be better defined as non-inflamed (group 1), mildly inflamed (group 2), and severely inflamed (group 3) than as TCD, CTRL, and UCD. This implies that inflammation status, rather than disease condition or Marsh score, is the main driver of this clustering (**Fig. 2A**). The non-inflamed cluster (group 1) consists of CTRLs and TCDs, as expected, whereas the mild inflammation cluster (group 2) is far more heterogeneous. The heterogeneity of group 2 could indicate that some CTRL individuals are clustered in group 2 due to ongoing non-CeD-associated inflammation, or that the TCD individuals in group 2 have not yet fully recovered or do not strictly adhere to GFD, and that the UCD patients in this group are in an early or non-severe phase of inflammation. In the severe inflammation group (group 3), only UCD cases are observed. Overall, this data suggests that both the UCD and TCD groups display heterogeneous transcriptomic features and that our cohort can better be classified based on the gene expression and inflammatory state of the EL.

To better identify the function of genes that contribute to the clustering, we analysed four clusters of genes in more detail (k=4, **Suppl. Fig. 5**). These genes have distinct roles in diverse biological pathways (**Fig. 2B**) (**Supplementary Table 3**). In general, cluster 1 genes, and to a lesser extent cluster 2 genes, are upregulated in UCD compared to the other conditions. In contrast, cluster 3 and 4 genes are predominantly downregulated in UCD. Cluster 4 gene upregulation seems to correlate best with CTRL and TCD samples. Cluster 1 is enriched for cell cycle and proliferation genes, including genes of the MCM protein complex, *CDC* genes, and polo-like kinase 1 (*PLK1*). Cluster 2 is enriched for genes associated with immune pathways and interleukin signalling, e.g. cathepsin G (*CTSG*), interleukin (*IL*) *21R*, *IL10*, C-X-C Motif Chemokine Ligand genes (e.g. *CXCL10* and *CXCL8*), and interferon gamma response genes including *IFNG*, signal transducer and activator of transcription 1 (*STAT1*), guanylate-binding protein genes (e.g. *GBP1*, *GBP5*, and *GBP4*), and suppressor of cytokine signalling 3 (*SOCS3*). Cluster 3 shows enrichment in genes that contribute to extracellular matrix organisation, including collagen genes (e.g. *COL1A2*, *COL3A1*, *COL18A1*, and *COL4A1*), matrix metallopeptidase genes (e.g. *MMP2* and *MMP9*) and laminin (e.g. *LAMA4*), and integrin genes (e.g. *ITGA9* and *ITGA5*), and to immunoregulatory interactions, including sialic acid–binding IG-like lectin genes (e.g. *SIGLEC1*, *SIGLEC7*, and *SIGLEC9*), transmembrane immune signalling adapter *TYROBP*, natural cytotoxicity triggering receptor 2 (*NCR2*), killer cell lectin-like receptor C1 (*KLRC1*), and CD40 ligand (*CD40LG)*. Genes in cluster 4 are enriched for digestion (e.g. guanylate cyclase activator 2A (*GUCA2A*), *GUCA2B*, sucrase-isomaltase (*SI*), *LCT*, and *TREH*), SLC-mediated transmembrane transport (*SLC* genes and lipocalin 15 (*LCN15*)) and laminin interaction pathways (e.g. *LAMA1*, *LAMA5*, *LAMB2*, *LAMB3*, and *COL7A1*). The changes in the pathways mentioned can be interpreted to be associated with specific CeD phenotypes. An increase in cell proliferation (cluster 1 genes) of both ECs and IELs might cause crypt hyperplasia and lymphocyte infiltration, whereas villus atrophy leads to disruption of digestion and absorption in the duodenum and could result from a problem in the basal lamina structure that contributes to a lack of mature enterocytes (as a result of downregulation of cluster 4 genes in UCD). Finally, CeD is associated with an increased type II interferon response (cluster 2 genes), causing a disruption of extracellular matrix and immunoregulatory interactions (cluster 3 genes).

### Differences in cell cycle, absorption, digestion, and basal lamina pathways confirm mucosal damage in UCD and TCD

Despite our finding that the overall transcriptome did not show a 1-to-1 relation with disease condition, we explored whether expression patterns of gene subsets can indicate disease status and predict the transcriptional heterogeneity observed in CTRL, UCD, and TCD. The first step is to consider the *APOA4:KI67* ratio as a proxy for villus health [23–25]. *APOA4* is a DE gene in cluster 4 and is known to be expressed in mature enterocytes [25], whereas *KI67* is a DE gene in cluster 1 and a marker of cell proliferation [25]. We observed that Marsh score correlated significantly with the *APOA4*:*KI67* ratio (**Suppl. Fig. 1**). Although the *APOA4*:*KI67* ratio changes do distinguish between UCD and CTRL/TCD, we could not distinguish TCD and CTRLs in our dataset using only this parameter (**Fig. 2C**, adjusted p-value < 0.01, Dunn test, Bonferroni correction), and thus it is not useful to set a clear threshold to differentiate between the severe, mild, or non-inflamed groups. As expected, DE genes found in clusters also correlated well with the *APOA4*:*KI67* ratio (**Fig. 2D**).

Next, we performed DE analysis on genes from the groups that appeared most variable: UCD severely inflamed (group 3) vs mildly inflamed (group 2), TCD mildly inflamed (group 2) vs non-inflamed (group 1), and UCD mildly inflamed vs TCD mildly inflamed (**Fig. 2E**, absolute L2FC > 1, adjusted p-value < 0.01) (**Supplementary Table 4**). This uncovered DE genes for each comparison, mostly between patient conditions for patients with different inflammation levels. However, when comparing mildly inflamed UCD vs TCD patients (group 2), we observed only a small number of DE genes, indicating that the transcriptome is similar between these disease conditions.

Most of the DE genes deregulated between UCD patients from inflammation group 3 vs 2 overlapped with DE genes of UCD vs TCD/CTRL (**Fig. 2F**) and had similar functions to cluster 1 and cluster 4 genes (**Suppl. Fig. 6**) (**Supplementary Table 5**): the upregulated DE gene set was enriched for cell cycle pathway genes and the downregulated gene set was enriched for ‘Digestion’ and plasma-lipoprotein-related pathways, indicating a decrease of functional enterocytes. These findings might suggest that UCD patients with mild inflammation are still in an early phase of active CeD, without full-blown damage to the EL.

For TCD individuals in the mild inflammation and non-inflamed groups, most DE genes overlapped with DE genes of UCD vs TCD/CTRL and to cluster 1 and 4 (**Fig. 2F**). Upregulated genes were enriched for cell cycle (e.g. *CDC* genes, *MCM10*, and *CDK1*), attributable to residual crypt hyperplasia or lymphocyte expansion [26–29], and downregulated genes were related to diverse transport mechanisms (e.g. *SLC* and *AQP* genes) and laminin interactions (e.g. *LAMB3*, *LAMC2*, and *LAMB2*) (**Suppl. Fig. 6**), which may be caused by a lack of mature enterocytes and a disruption of the basal lamina of the EL, respectively. Our results suggest that within the TCD group there are patients still in recovery whose gene profile contains DE genes previously identified in gene cluster 1 and 4. The TCD individuals in the non-inflamed group were similar to controls, indicating that these subjects had fully recovered after initiation of a GFD. Both the TCD and UCD subjects in the mildly inflamed group were similar, which could indicate partial recovery after GFD or mild damage due to non-gluten-related inflammatory response.

Overall, gene expression variation indicates different levels of inflammation in the EL, suggesting different levels of intestinal dysfunction in individuals, as indicated by the deregulation of genes found in cluster 1, related to cell cycle, and in cluster 4, associated with decreased absorptive and metabolic function, regardless of Marsh scores and disease condition. The transcriptomic state of the EL is thus a good indicator of the inflammatory state and may prove helpful to further classify CeD patients as non-inflamed, mildly inflamed, or severely inflamed. However, the transcriptomic data cannot predict whether mildly inflamed tissue indicators are caused by CeD onset (or gluten-induced damage before villus atrophy) or by inflammation induced by another non-gluten factor.

### Immune- and extracellular-matrix-associated genes classify untreated and treated CeD patients

While the genes present in clusters 1 and 4 distinguish between various states of inflammation and dysfunction of the small intestine, they do not clearly distinguish CTRLs from TCDs and TCDs from UCDs. To assess whether we could predict disease condition based on gene expression, despite the heterogeneity in the inflammatory state of patients, we applied a rank-based single-sample gene set scoring method using clusters of DE genes (**Fig. 3**) (**Supplementary Table 6**) [30]. Ideally, this method should result in a clear separation of samples in which active CeD cases obtain positive scores and CTRLs obtain negative scores (or vice versa). The latest is achieved by using upregulated and downregulated DE genes as inputs. We assessed distinctions between CTRL, TCD, and UCD patients based on singscores using all DE genes or combinations of clusters of DE genes (**Suppl. Fig. 7**, adjusted p-value < 0.01, Dunn test, Bonferroni correction). We found that the patient groups could be best distinguished based on only cluster 2 and 3 genes as this yielded the lowest adjusted p-values (UCD vs CTRL (p-value = 1.29×10^-15^) and TCD vs CTRL (p-value = 1.88×10^-03^)), despite using a lower total number of genes (**Fig. 3A**). Moreover, cluster 2 and 3 together showed a clear separation of UCD (median = 0.28, IQR = 0.23–0.34) and CTRL (median = −0.29, IQR = −0.36–-0.21) based on singscore, a separation that was not observed when taking only the *APOA4*:*KI67* ratio into consideration. In contrast, TCD samples scored in between the other conditions (median = −0.08, IQR = −0.15–0).

**Figure 3.**
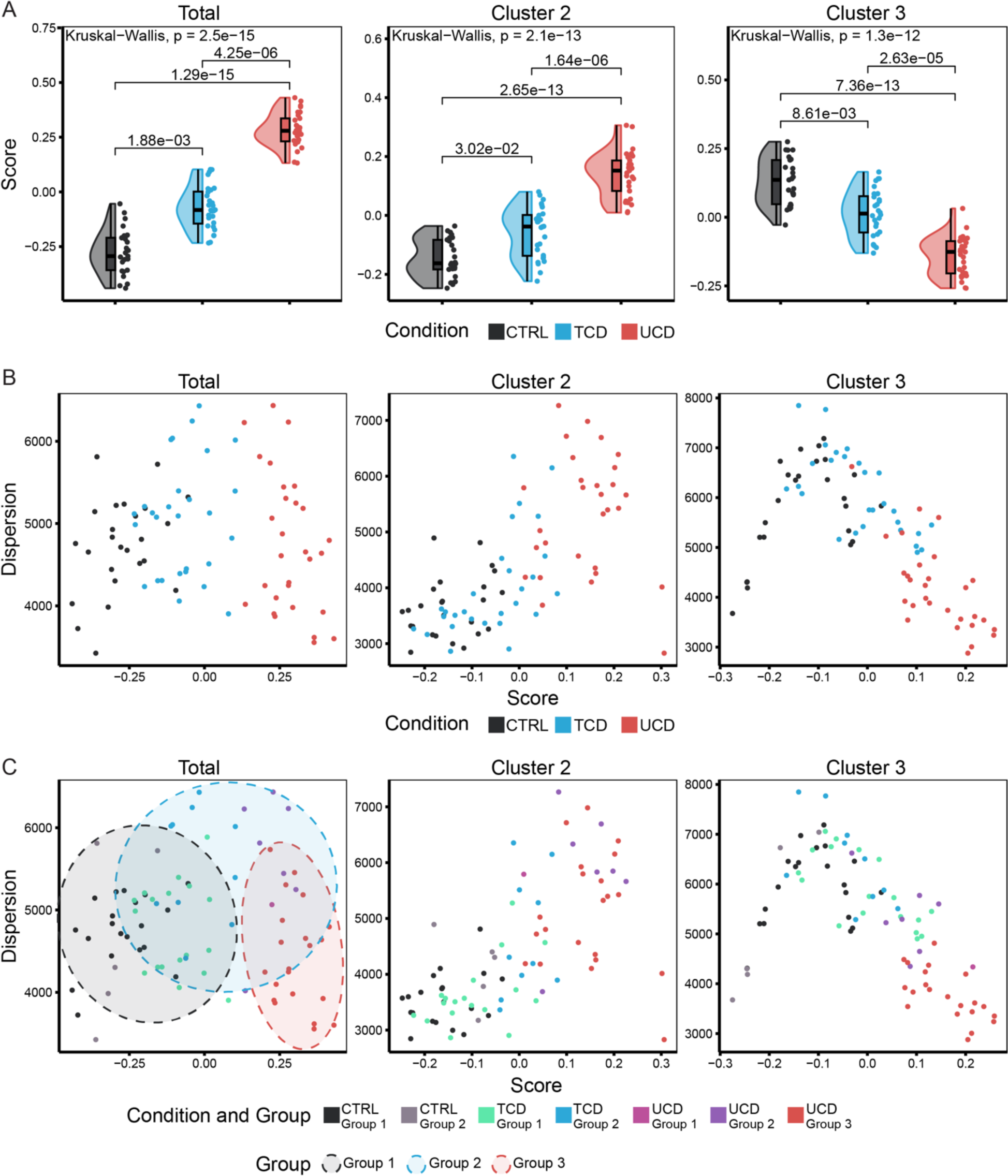
**DE gene clusters 2 and 3 help identify CeD status**. **(A)** Singscore (y-axis) of samples when using gene expression of clusters 2 and 3 (left), cluster 2 (centre), or cluster 3 (right), divided and coloured by CeD condition (x-axis) (Kruskal-Wallis test and Dunn test as post-hoc test). **(B & C)** Dot plots of singscore (x-axis) versus dispersion (y-axis). In (B), samples are coloured by CeD condition. In (C), samples are coloured by condition and group. Ellipses show groupings indicated by multivariate t-distribution.

Finally, to further characterise the heterogeneity in our samples, we assessed their relative rank dispersion (**Fig. 3B**). If samples have a similar score but a different rank dispersion, it may indirectly reflect a different regulation of the gene sets used [30], i.e. these genes may change expression in the same direction but at different levels. Indeed, the mildly inflamed group displays higher dispersion, indicative of larger variation within this group, whereas the severely inflamed and non-inflamed groups tend to display lower dispersion. To validate the ability of cluster 2 and 3 genes to define disease state, we used the genes in these clusters to assign disease conditions using a replication transcriptomic dataset generated from 51 duodenal biopsies from CeD patients and 44 healthy controls described in literature [31]. Again, we found the best predictors when using either cluster 2 and 3 together or when using all four clusters (43/44 individuals were predicted as CTRL and 43/51 as UCD subjects, using either approach). Other cluster combinations performed less well (**Suppl. Fig. 8**). We thus conclude that the genes in clusters 2 and 3, which mainly encompass immune and extracellular matrix genes, have the potential to distinguish between UCD, TCD, and CTRL. Moreover, as genes from clusters 2 and 3 are already deregulated in mild inflammation UCD individuals, with immune genes upregulated and extracellular matrix genes downregulated, we propose that these genes are early responders in CeD inflammation.

### Deconvolution of eQTL analysis pinpoints genes affected by genetics in the immune and epithelial cell compartment

As patient genetics is likely to contribute to the heterogeneity between patients, we evaluated how single nucleotide polymorphisms (SNPs) associated with CeD by genome-wide association studies affect the expression of the genes in the associated genetic loci [32–34]. To minimise the multiple-testing burden, we performed *cis*-eQTL analysis using the lead SNPs in each CeD-associated locus, including only genes located within 250 kb of the lead SNP [34,35]. Despite minimising the multiple-testing burden, most of the bulk and cell-type-mediated *cis*-eQTLs detected are only suggestive, likely due to power limitations given the low sample numbers. Nonetheless, we uncovered 25 eQTL genes with a suggestive significant effect (p-value < 0.005) when analysing bulk-RNA results obtained from the EL (**Fig.**) (**Supplementary Table 7**). The top 5 eQTL genes were zinc finger protein 57 (*ZFP57*), two HLA genes (*HLA-G* and *HLA-K*), membrane-metalloendopeptidase-like 1 (*MMEL1*), and *IL18R1*. The first three genes are in the HLA locus on chromosome 6. The other two are located on chromosome 1 and 2, respectively. The *MMEL1* and *IL18R1* SNP-gene pairs were previously reported by the eQTLGen Consortium, which used whole-blood RNA [36], indicating that these eQTLs are not specific to duodenal tissue. We also deconvoluted eQTL effects in both major cell types in our samples: epithelial and immune cells. For this, we applied Decon-QTL [37], which imputes cell-type-mediated eQTLs using known cell proportions (**Supplementary Table 8**). In total, we observed 6 EC and 15 immune cell eQTL genes, of which 3 and 6 genes, respectively, overlapped with those obtained in the bulk tissue eQTL analysis (**Fig. 4A**, p-value < 0.01). Of these, *IL18R1* and *IL18RAP* were eQTL genes in both bulk tissue and immune cells, but not in ECs, with a clear immune-related function (**Fig. 4B,C**). Neither of these genes were DE however, indicating that genetic effects may function independently of variation in gene expression, even though both contribute to disease state. Additionally, we observed immune-cell-mediated eQTL genes, e.g. membrane spanning 4-domains A14 (*MS4A14*), which is expressed in myeloid cells [38], and T cell activation RhoGTPase activating protein (*TAGAP*), which has a role in both T cells and myeloid cells [39]. The functions of the remaining eQTL genes specific to ECs and immune cells are less clear (**Supplementary Table 7**).

**Figure 4.**
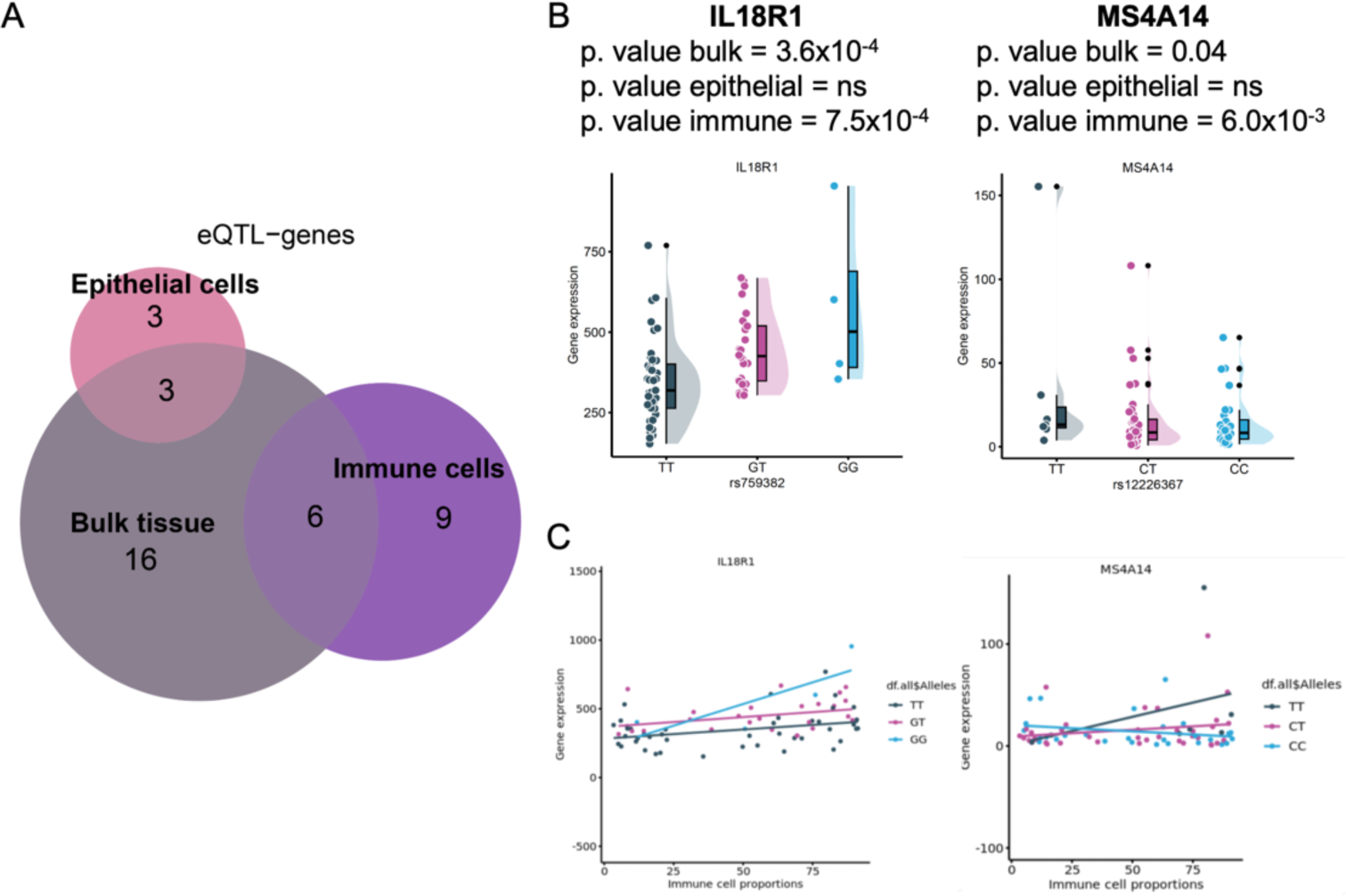
Bulk and cell-type-mediated eQTLs in CeD context. **(A)** Overlap of eQTL genes represented as Euler diagrams classified as bulk tissue, epithelial cell, and immune cell eQTLs. Examples of eQTLs visualised as dot plots of gene expression (y-axis) versus **(B)** genotype (x-axis) and **(C)** proportions of immune cells (x-axis). Samples coloured by genotype.

## Discussion

To assess the heterogeneity of the duodenal EL in CeD, we characterised the transcriptomic landscape of duodenal EL in treated and active CeD transcriptional profiles. We defined three groups based on the inflammation status (non-inflamed, mild inflammation, and severe inflammation) that were correlated with but not specific to disease state. UCD patients were found to display mild or severe inflammatory transcriptional features, whereas TCD patients exhibited a transcriptional phenotype similar to that of CTRL individuals or to UCD patients with mild inflammation. Two clusters of genes enriched for immune and extracellular matrix and barrier function (**Fig. 5**) yielded the best classification into specific conditions (CTRL, TCD, and UCD).

**Figure 5.**
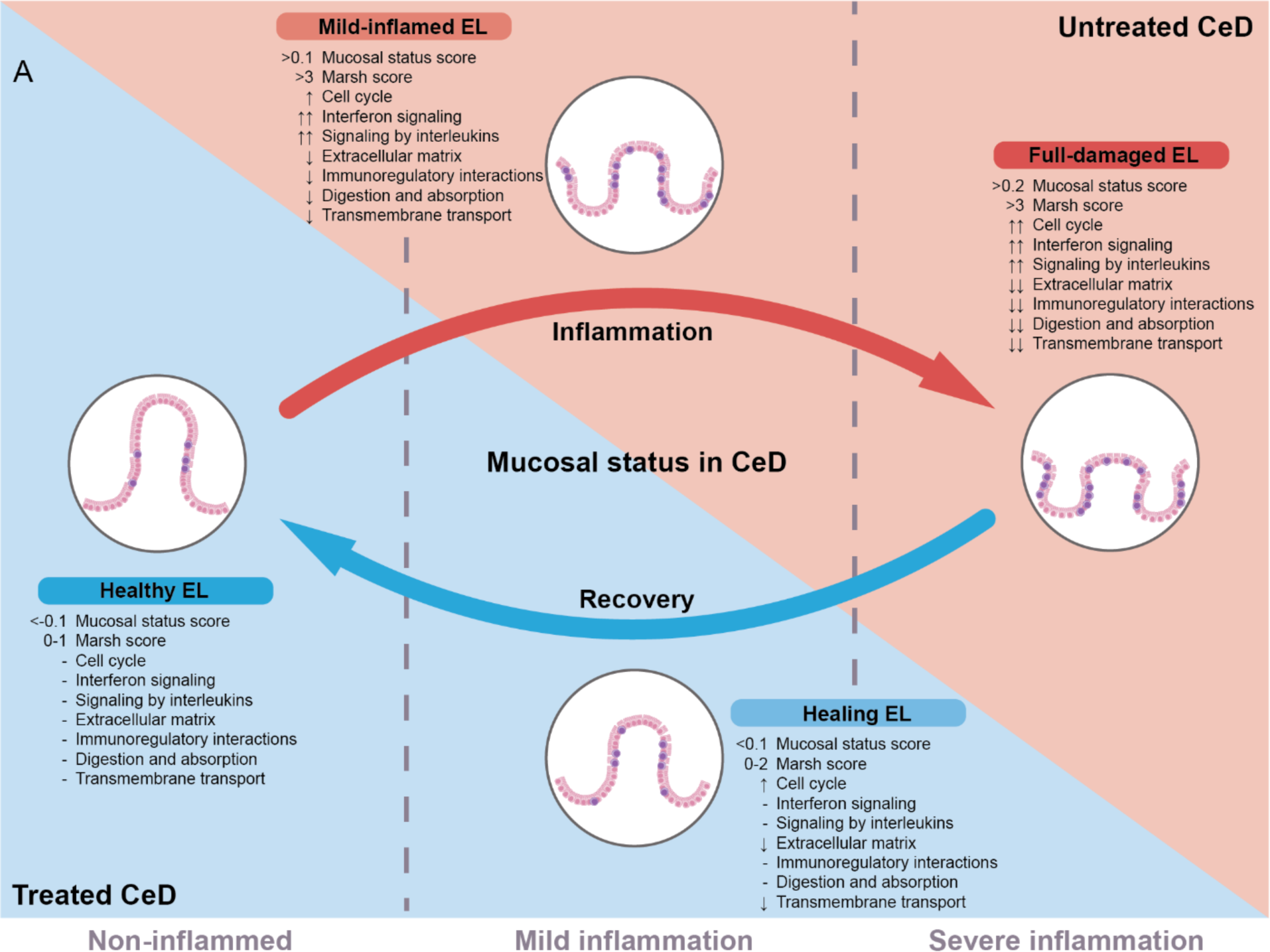
Interpretation of the transcriptomic results associated with the mucosal status of the duodenal EL in CeD. In untreated CeD (UCD), two distinct states are observed: mildly and severely inflamed EL. Mild-inflamed EL already shows a decreased APOA4:KI67 ratio, suggesting a decreasing villus height. Interleukin signalling pathways are already upregulated. Although the cell cycle is upregulated in mild inflammation, it is not as pronounced as in severely inflamed EL. Moreover, extracellular matrix and digestive functions are disrupted in mild inflammation. Full damage EL displays extreme phenotypes: upregulation of pro-inflammatory immune pathways; downregulation of extracellular matrix, membrane transport and metabolic pathways; strong upregulation of the cell cycle (stronger than in mild-inflamed EL); and strong downregulation of digestive and absorptive processes (stronger than mild-inflamed EL). In UCD, the APOA4:KI67 ratio is lowest, suggesting severe villus atrophy. In TCD, the EL is recovering or already fully recovered. The TCD transcriptome of mildly inflamed EL is similar to that in UCD with the same inflammation status. However, TCD Marsh scores of 0–2 are lower than those of UCD (3), indicating less epithelial damage. Moreover, the gene expression singscore suggests that the degree of damage (cluster 3 of DE genes) and inflammation (cluster 2 of DE genes) is lower than in UCD. Healthy EL has characteristics similar to CTRL EL, with similar APOA4:KI67 ratios and Marsh scores ≤ 1, indicating a healthy phenotype. Arrows depict the direction of pathway deregulation. “-” signifies normal expression levels.

Determination of DE genes from duodenal EL can help uncover the genes and pathways that are involved in the tissue damage associated with CeD and shed light on the causes of the phenotypical heterogeneity in the disease. Our data uncovered genes explaining inflammation status (and presumably villous atrophy) and disease severity. UCD cases are characterised by upregulation of cell cycle and proliferation genes (e.g. *MCM* and *CDC* genes, *PLK1*, and *CLK1*) and downregulation of digestive (e.g. *GUCA2A*, *GUCA2B*, *SI*, *LCT*, and *TREH*), transmembrane transport (*SLC* genes), and basal lamina genes (e.g. *LAMA1*, *LAMA5*, *LAMB2*, *LAMB3*, and *COL7A1*) (**Fig. 2**). DE genes were further classified into four clusters, leading us to distinguish cluster 1 as enriched for cell proliferation functions and cluster 4 as enriched for digestion, absorption, and laminin functions. These pathways are mainly deregulated in UCD and are indicative of an expanding IEL population and EC proliferation that in turn may result in crypt hyperplasia [26,27,29]. The downregulation of digestive, transport, and laminin interaction pathways suggest a loss of intestinal epithelial functions, perhaps due to the loss of enterocytes due to villous atrophy. Downregulation of transport-related genes is in line with previous observations by Laforenza et al, who found alteration of expression of aquaporin genes (*AQP7, AQP10*, and *AQP11*) and solute carrier genes (*SLC5A1* and *SLC15A1*) in the same direction [41]. Similarly, in intestinal biopsies of CeD patients, Veberke et al found a weaker staining of proteins related to extracellular matrix, such as collagen IV, laminin and fibronectin, indicating a concordance with the trends we observe in our study [42]. In our TCD cases, upregulation of cell cycle and proliferation genes and downregulation of digestive, transmembrane transport, and basal lamina genes distinguishes between the non-inflamed and mildly inflamed states within TCD, pointing to incomplete restoration of the EL and enterocyte function. Finally, cluster 1 (cell cycle and proliferation genes) and cluster 4 (digestive, absorptive, and basal lamina genes) genes recapitulate the mucosal inflammation status of EL in CeD, making it possible to identify mild or severe inflammation in untreated CeD cases and non-inflamed or mild inflammation in treated CeD cases.

Leonard et al [43] also studied the duodenal transcriptome of UCD, TCD patients, and CTRLs and observed that most DE genes (89%) corresponded to the comparisons of active CeD biopsies versus any other condition, identifying DE genes and enriched pathways similar to those we observe: genes associated to CeD (*IFNG, CDC45,* and *MCM*2), interleukins (*IL10* and *IL17A*), chemokines (*CXCL3, CXCL9, CXCL10,* and *CXCL11*), and cell adhesion molecules (*CLDN18*). However, when comparing TCD cases versus controls they identified more DE genes (290 vs 30 DE genes). We speculate that this divergence may be attributed to differences between the processing/preparation of the samples or the cell composition of the tissue used, i.e. EL versus whole duodenal biopsy, where the latter also includes the lamina propria. Lamina propria could better recapitulate the residual immune dysregulation after GFD in CeD cases. Indeed, immune-associated genes were not enriched in our DE analysis within TCD (mild inflammation vs non-inflamed) and within UCD (severe vs mild inflammation), supporting the idea that lamina propria may be a better context to observe the immune response before and after GFD [44–46].

The DE genes in clusters 2 and 3 exhibited up- vs downregulation, respectively, in UCD cases, regardless of disease severity. Indeed, these genes were sufficient to distinguish UCDs from CTRLs and TCD, as shown by the singscore results (**Fig. 3**). Therefore, the genes in clusters 2 and 3 might be classified as genes involved in CeD onset and are deregulated independently of the loss of intestinal function and increased proliferation. Indeed, cluster 2 genes are enriched in immune-related pathways, such as signalling by interferon gamma (e.g. *IFNG*, *STAT1*, and *GBP1*) and by interleukins (e.g. *IL21R*, *IL17A*, and *IL10*), and include some of the top genes found in CeD-associated IELs (e.g. *KLRC2, RTKN2, BUB1B, TNFRSF9,* and *CISH)* [16,17,22].

Moreover, some cluster 2 genes, including *IFNG*, *GBP5*, *CXCL10*, and *UBD,* have been documented to be upregulated in active CeD even prior to diagnosis, as reported by Bradge et al [47]. Finally, the singscores in TCD fall between those of CTRLs and UCD, which may suggest that the duodenal mucosa is still not completely recovered. Suboptimal recovery of the villi might be consistent with an observation by others that CeD-associated IEL populations remain in the EL even after start of a GFD [17,22]. Cluster 3 is enriched for genes involved in extracellular matrix organisation (e.g. *COL23A1*, *MMP2*, and *DCN*) and immunoregulatory interactions (e.g. *TYROBP*, *SIGLEC1*, and *CD40LG*), among others. Cluster 3 also contains downregulated genes related to the inflammatory process in CeD, and some of these downregulated immune-related genes (e.g. *GNLY*, *KLRC1*, *TYROBP, FCER1G, GZMK,* and *SH2D1B*) are implicated in natural IEL function in a healthy mucosal environment [17]. Cluster 2 and 3 genes clearly distinguished UCD from TCD and CTRLs, whereas genes in cluster 1 and 4 were more indicative of villous atrophy and tissue damage in UCD or incomplete recovery in TCD (**Suppl. Fig. 7**). The cluster 2 and 3 genes together were also more predictive of disease status than the often-used *APOA4:KI67* ratio, suggesting they represent a novel means to capture mucosal status and immune processes and predict IEL populations in CeD development.

Our eQTL and deconvolution analysis resulted in more immune-cell-mediated hits than EC-mediated hits. *IL18R1*:rs759382 is both a bulk eQTL and an immune-mediated eQTL, and it was reported previously as a blood eQTL [36]. Although *IL18R* is not DE, its ligand, *IL18*, was downregulated in UCD and present in cluster 4. IL18R is found on the surface of several cell types, including ECs, dendritic cells, and subsets of lymphocytes, and thus its interaction with IL18 may be altered as a consequence of genetic associations in the context of CeD. IL18 and IL18R have pleiotropic functions in maintaining inflammation in CeD [48,49] and affect epithelial barrier function in colitis [50]. The eQTL *MS4A14*:rs12226367 was only found in the immune-compartment-mediated eQTL analysis. *MS4A14* is expressed mainly in monocytes and myeloid cells and is located in the same locus as other members of the MS4A family [38]. Although to our knowledge this is a novel eQTL effect, rs12226367 was previously associated to the expression of *MS4A6A* [51]. Overall, we obtained a relatively small number of eQTL genes, likely due to the lack of power because of the limited size of our cohort. Moreover, to standardise the eQTL analysis and eliminate the influence of CeD status, we controlled for CeD condition without considering inflammation status. This approach may have hampered the analysis of genetic effects on gene expression in our data. Conducting additional eQTL analyses in larger and more precisely stratified cohorts for CeD, and incorporating factors such as inflammation status, could potentially address this limitation. This would enable a more thorough exploration of the interactions between cell-type-mediated eQTL effects and CeD condition.

In conclusion, using cell cycle, digestive, absorptive, and basal lamina genes, we can stratify CeD patients based on their inflammation/villus damage status. Remarkably, despite heterogeneity in active and treated CeD cases, we were able to accurately classify untreated CeD patients, treated CeD patients, and controls based on immune and extracellular matrix genes, which we suggest play an important role in CeD pathophysiology. Overall, we identified genes expressed in the duodenal EL whose expression might be used as biomarkers to assess CeD condition and its mucosal and immune status.

## Methods

### Ethical considerations and study design

CeD patients were included at the Rikshospital in Oslo, Norway. Briefly, written informed consent was given by all patients, who donated blood before going through an upper endoscopy to obtain small intestine biopsies for diagnosis and research purposes. All methods were performed in accordance with relevant guidelines and regulations.

A total of 113 participants were included in this study and classified into three different groups: controls (CTRL), treated CeD patients (TCD), and untreated CeD patients (UCD). Controls (n=40) consisted of volunteers undergoing upper endoscopy for complaints unrelated to CeD. TCD (n=39) were CeD patients on a GFD with a Marsh score ≤ 2. UCD (n=34) were CeD patients previously diagnosed or suspected of having CeD with a Marsh score of 3.

### Genotyping and quality control

DNA was isolated from blood samples and genotyped using Infinium Global Screening Array-24v1.0. Standard quality control (QC) procedures were used to remove low quality variant calls. Genotypes were imputed with the Michigan Imputation Server using the Haplotype Reference Consortium panel v1.172, as described previously [52]. SNPs with an imputation score < 0.8, a Hardy-Weinberg equilibrium p-value < 1×10^−4^, a call rate < 95%, or a minor allele frequency < 0.1 were filtered out.

### Preparation of small intestine biopsies

CeD patients were included at the Rikshospital, Oslo. Biopsies were obtained by upper endoscopy and incubated under agitation in PBS+EDTA solution on ice for 5 minutes. Cells were collected from the supernatant by pelleting at 300 g for 5 min and passed through a 70 μm cell strainer. A fraction of these cells was analysed by FACS, quantified by trypan blue exclusion medium. The rest was pelleted and resuspended in Lysis/Binding medium (AM1560). Lysed cells were frozen and stored until RNA extraction.

### FACS preparation and analysis

Single-cell suspensions were stained with the antibody panel depicted in **Supplementary Table 8**. Briefly, cells were centrifuged at 300 g for 5 min and then resuspended in 100 μL PBS supplemented with 2% FCS. Next, cells were stained for 30 min at 4℃, after which they were washed twice and resuspended in 400 μL PBS supplemented with 2% FCS. FACS data was generated using the BD LSR-II system (BD Bioscience) and analysed using FlowJo v10.

### RNA isolation of EL

Samples were processed and sequenced in two sequencing batches. Total RNA was extracted from 113 samples of the EL cell fraction using the mirVana™ miRNA Isolation Kit (AM1560). Up to 12 samples were processed simultaneously, ending with 10 batches of RNA extraction. The RNA was quantified, and integrity was confirmed by different approaches: Nanodrop spectrometry, Qubit RNA BR Assay Kit (Q10210), Qubit RNA HS Assay Kit (Q32852), Agilent RNA ScreenTape (5067-5576), and Agilent High Sensitivity RNA ScreenTape (5067-5579). Before library preparation, RNA integrity (RIN) and concentration was measured again and used to calculate a ratio of degradation ([RNA concentration after extraction]/[RNA concentration before library preparation]). Samples with a confirmed RIN > 6 and a concentration > 0.5 ng/µL were sequenced, resulting in a total of 90 samples.

### RNA library preparation

RNA library preparation and sequencing was performed at GenomeScan B.V., Plesmanlaan 1d, 2333 BZ, Leiden. Sample preparation was performed according to the protocol “NEBNext Ultra II Directional RNA Library Prep Kit for Illumina” (NEB #E7760S/L). Briefly, mRNA was isolated from total RNA using oligo-dT magnetic beads. After fragmentation of the mRNA, a cDNA synthesis was performed. This was used for ligation with the sequencing adapters and PCR amplification of the resulting product. The quality and yield after sample preparation was measured with the Fragment Analyzer. The size of the resulting products was consistent with the expected size distribution (a broad peak between 300–500 bp). NovaSeq6000 was used for clustering and DNA sequencing following manufacturer’s protocols using a concentration of 1.1 nM DNA. Image analysis, base calling, and QC were performed with the Illumina data analysis pipeline RTA (version 3.4.4) and Bcl2fastq (version 2.20).

### RNA-seq quantification and QC

The adapters for sequencing were trimmed from fastQ files and aligned to build human_g1k_v37 ensembleRelease 75 reference genome using Hisat (version 0.1.5) [53] with default settings. SAMtools (version 1.2) [54] was used to sort the aligned reads before gene quantification. Gene-level quantification was performed by HTSeq-count HTSeq (version 0.6.1p1) [55] using -- mode=union. QC metrics were calculated for the raw sequencing data using the tool FastQC (version 0.11.3) [56]. QC metrics were calculated for the aligned reads using Picard-tools (version 1.130) [57].

The raw count matrix, containing 53,042 transcripts and 90 samples, was first filtered to remove zero-variance and non-expressed genes by selecting only genes that had at least 10 reads in 10 samples. This resulted in 20,498 genes for further analysis. Next, to remove sample outliers, we explored the first four PCs using the 1000 most variable genes, experimental variables, and sequencing QC metrics. In total, we removed eight outliers that showed at least three of the following criteria: outlier in the PC analysis, unique pair read percentages < 10%, unmapped read percentage > 10%, RNA concentration < 1.5 ng/µL, total number of reads > 3 million, sample present in batch 9 of RNA extraction, or ratio of degradation > 10 (calculated as [RNA concentration after extraction]/[RNA concentration before library preparation]). Y chromosome genes were also excluded from the dataset. The final dataset consisted of 82 samples and 20,468 genes.

### DE analysis

The DE effects of different conditions were quantified using the R package DEseq2 (version 1.34.0) [58], including sex, age, sequencing batch, total reads, RNA integrity, and %GC content as covariates in the DE model. DE effects were calculated by comparing UCD vs CTRL, TCD vs CTRL, and UCD vs TCD. DE effects were filtered on having an absolute L2FC ≥ 1 and an adjusted p-value < 0.01. The remaining DE genes were used for interpretation and downstream analysis.

### Clustering of DE genes and sample groups

DE genes were clustered into groups as follows. The gene expression matrix was VST-normalised using DESeq2 (version 1.34.0) [58]. The matrix was then filtered to include only DE genes. Next, gene expression was centred to mean 0 and scaled. DE genes were clustered using k-means clustering (k=4) on a Euclidean distance matrix using the R package ComplexHeatmap (version 2.10.0) [59]. Samples were also clustered following a similar approach to that used for DE genes, but with k=3. The cluster number (k) was determined by comparing three different methods to obtain the optimal number of clusters that are biologically informative. These methods were obtaining gap statistic (500 permutations), average silhouette width, and total within sum-of-square when clustering from 1 to 10 groups. For this analysis, we used the R packages cluster (version 2.1.6) [60] and factoextra (version 1.0.6) [61].

### Pathway enrichment analysis

Reactome pathways [62] were used to identify the pathways or biological processes that were enriched for each set of genes. This analysis was performed using the R package clusterProfiler (version 3.14.3) [63]. P-values were adjusted using the Benjamin-Hochberg procedure to account for multiple testing.

### APOA4:KI67 ratio calculation

The gene expression matrix was VST-normalised using DESeq2 (version 1.34.0) [58]. The APOA4:MKI67 ratio was calculated by dividing the gene expression of *APOA4* by that of *MKI67* for each sample.

### Scoring of mucosal status in samples

The mucosal status score for each sample was calculated using the methodology implemented in the R package singscore (version 1.14) and the pipeline recommended by authors [30]. Briefly, singscore is a single-sample gene set scoring method that values individual samples without relying on other samples included in the dataset. It can use sets of genes that are up- and downregulated to score molecular phenotypes. First, the RNA matrix is ranked using the rankGenes() function. This rank matrix, along with the gene sets, is then passed to the simpleScore() function, which returns the output with scores and dispersions. As gene sets, the input included all possible combinations of DE genes included for each previously identified cluster. Since we know the direction of DE genes, we used clusters 1 and 2 as upregulated sets and clusters 3 and 4 as downregulated sets. Resulting scores ranged between −1 and 1. Values > 0 indicate an enrichment of the set of genes used as input, whereas a negative value corresponds to a depletion of genes. When only one cluster is used (i.e. only cluster 3), it is passed on to the upSet argument, which uses the cluster as an upregulated gene set. Additionally, the dispersion of scores was measured using the median absolute deviation of the gene set ranks, as described by the authors.

### Bulk eQTL analysis

For bulk eQTL mapping, we tested for effects between genes and CeD-associated SNPs [32,34] located within 250 kb of a gene centre. The RNA matrix was TMM-normalised using R package edgeR (version 3.36.0) [64] and corrected for covariates (sex, age, sequencing batch, total reads, RNA integrity, %GC content, Marsh scores, condition, and APOA4:KI67 ratio), the first four PCs derived from gene expression, and four multidimensional scaling components derived from the genotype data. eQTLs were declared to be suggestive at FDR < 0.01. QTL mapping was performed using an eQTL pipeline that was described previously [65].

### Deconvolution of eQTL effects in cell types

For this analysis, we employed the method Decon-QTL [37], testing for the same effects as in the bulk eQTL mapping. Gene counts were log2 transformed and corrected for sex, age, sequencing batch, total reads, RNA integrity, %GC content, Marsh scores, condition, APOA4:KI67 ratio, the first four PCs derived from gene expression and four multidimensional scaling components derived from the genotype data. The corrected expression data was then exponentiated to maintain the original linear relationship across read counts and cell proportions. As cell counts, we used proportions of major immune and epithelial cells. Cell-type-mediated QTLs were considered suggestive at a p-value < 0.01

### Statistical methods

Statistical analyses were performed in R (version 3.6.3) [66], unless otherwise specified. Visualisation of results was done using the R package ggplot2 (version 3.3.0) [67].

## Acknowledgements

We thank all patients that have donated biological material to this study. We thank Kate Mc Intyre for editing the manuscript. A.D.R.S. is supported by a CONACYT-I2T2 scholarship (no. 459192). S.Z. was supported by the K.G. Jebsen foundation. S.W. is supported by the Netherlands Organ-on-Chip Initiative, an Netherlands Organization for Scientific Research (NWO) Gravitation project (024.003.001) funded by the Ministry of Education, Culture and Science of the government of the Netherlands. I.H.J. is supported by a Rosalind Franklin Fellowship from the University of Groningen and am NWO VIDI grant (no. 016.171.047). L.F. is supported by a grant from the Dutch Research Council (grant no. ZonMW-VICI 09150182010019), a European Research Council Starting Grant (grant agreement 637640 (ImmRisk)), an Oncode Senior Investigator grant, and sponsored research collaborations with Biogen, Roche, and Takeda. L.F. also received support from the European Union’s Horizon Europe Research and Innovation Programme under grant agreement No 101057553.

## Author contributions

A.D.R.S. and S.Z. performed wet-lab experiments. K.E.A.L. recollected samples and metadata.

A.D.R.S. and S.Z. conceived and wrote the manuscript. A.D.R.S. performed the data analyses.

R.A.G. assisted in the application of the deconvolution eQTL method. M.V. and L.F. supported in the implementation of the bulk eQTL method. I.H.J., K.E.A.L., and S.W. supervised and edited the manuscript.

## Data availability

The RNA-seq count tables and FACS counts required to reproduce this study are provided as supplementary files. The raw RNA-seq and genotypes supporting this study are available upon request to the authors as this data is privacy-sensitive. All code and scripts used to generate the results and figures are available on Github (https://github.com/aarondrs2/CeD_EpithelialLining_RNAseq_2024).

## Ethics statement

Biological material was obtained from patients according to protocols approved by the regional ethics committees (University Medical Center and University of Oslo), and the individuals donating material gave their written informed consent.

## Supplementary Material

### DATA

**Supplementary data 1.** Matrix of RNA-seq raw counts of genes per sample.

### TABLES

**Supplementary table 1.** Sample sheet and QC details

**Supplementary table 2.** List of DEGs: comparison between disease conditions

**Supplementary table 3.** Enriched pathways of DEGs: comparison between disease conditions

**Supplementary table 4.** List of DEGs: comparison between inflammation groups

**Supplementary table 5.** Enriched pathways of DEGs: comparison between inflammation groups

**Supplementary table 6.** Singscores of samples using DEGs

**Supplementary table 7.** List of bulk and predicted-cell-type eQTLs

**Supplementary table 8.** Cell-type counts by FACS analysis and antibody panel

## References

[1] Al-Toma, A., Volta, U., Auricchio, R., Castillejo, G., Sanders, D.S., Cellier, C. et al. (2019) European Society for the Study of Coeliac Disease (ESsCD) guideline for coeliac disease and other gluten-related disorders. United European Gastroenterology Journal, 7, 583–613. 10.1177/2050640619844125

[2] Mowat, A.M. and Agace, W.W. (2014) Regional specialization within the intestinal immune system. Nature Reviews Immunology, 14, 667–85. 10.1038/nri3738

[3] Iversen, R. and Sollid, L.M. (2023) The Immunobiology and Pathogenesis of Celiac Disease. Annual Review of Pathology: Mechanisms of Disease, 18, 47–70. 10.1146/annurev-pathmechdis-031521-032634

[4] Mayassi, T. and Jabri, B. (2018) Human intraepithelial lymphocytes. Mucosal Immunology, 1–9. 10.1038/s41385-018-0016-5

[5] Arentz-Hansen, H., Körner, R., Molberg, Ø., Quarsten, H., Vader, W., Kooy, Y.M.C. et al. (2000) The intestinal T cell response to α-gliadin in adult celiac disease is focused on a single deamidated glutamine targeted by tissue transglutaminase. Journal of Experimental Medicine, 191, 603–12. 10.1084/jem.191.4.603

[6] Simon-Vecsei, Z., Király, R., Bagossi, P., Tóth, B., Dahlbom, I., Caja, S. et al. (2012) A single conformational transglutaminase 2 epitope contributed by three domains is critical for celiac antibody binding and effects. Proceedings of the National Academy of Sciences of the United States of America, 109, 431–6. 10.1073/pnas.1107811108

[7] Fleckenstein, B., Qiao, S.W., Larsen, M.R., Jung, G., Roepstorff, P. and Sollid, L.M. (2004) Molecular Characterization of Covalent Complexes between Tissue Transglutaminase and Gliadin Peptides. Journal of Biological Chemistry, 279, 17607–16. 10.1074/jbc.M310198200

[8] Stamnaes, J. and Sollid, L.M. (2015) Celiac disease: Autoimmunity in response to food antigen. *Seminars in Immunology*, Elsevier Ltd. 27, 343–52. 10.1016/j.smim.2015.11.001

[9] Molberg, Kett, K., Scott, H., Thorsby, E., Sollid, L.M. and Lundin, K.E.A. (1997) Gliadin specific, HLA DQ2-restricted T cells are commonly found in small intestinal biopsies from coeliac disease patients, but not from controls. Scandinavian Journal of Immunology, 46, 103–8. 10.1046/j.1365-3083.1997.d01-93.x-i2

[10] Mesin, L., Sollid, L.M. and Niro, R. Di. (2012) The intestinal B-cell response in celiac disease. Frontiers in Immunology, 3, 1–12. 10.3389/fimmu.2012.00313

[11] Cerf-Bensussan, N., Guy-Grand, D. and Griscelli, C. (1985) Intraepithelial lymphocytes of human gut: isolation, characterisation and study of natural killer activity. Gut, 26, 81–8. 10.1136/gut.26.1.81

[12] Sánchez-Castañon, M., Castro, B.G., Toca, M., Santacruz, C., Arias-Loste, M., Iruzubieta, P. et al. (2016) Intraepithelial lymphocytes subsets in different forms of celiac disease. Autoimmunity Highlights, 7. 10.1007/s13317-016-0085-y

[13] Jabri, B., De Serre, N.P.M., Cellier, C., Evans, K., Gache, C., Carvalho, C., et al. (2000) Selective expansion of intraepithelial lymphocytes expressing the HLA-E-specific natural killer receptor CD94 in celiac disease. Gastroenterology, 118, 867–79. 10.1016/S0016-5085(00)70173-9

[14] Abadie, V., Discepolo, V. and Jabri, B. (2012) Intraepithelial lymphocytes in celiac disease immunopathology. Seminars in Immunopathology, 34, 551–66. 10.1007/s00281-012-0316-x

[15] Carrasco, A., Mañe, J., Santaolalla, R., Pedrosa, E., Mallolas, J., Lorén, V. et al. (2013) Comparison of lymphocyte isolation methods for endoscopic biopsy specimens from the colonic mucosa. Journal of Immunological Methods, 389, 29–37. 10.1016/j.jim.2012.12.006

[16] Kornberg, A., Botella, T., Moon, C.S., Rao, S., Gelbs, J., Cheng, L. et al. (2023) Gluten induces rapid reprogramming of natural memory αβ and γδ intraepithelial T cells to induce cytotoxicity in celiac disease. Science Immunology, 8, 1–29. 10.1126/sciimmunol.adf4312

[17] Atlasy, N., Bujko, A., Bækkevold, E.S., Brazda, P., Janssen-Megens, E., Lundin, K.E.A. et al. (2022) Single cell transcriptomic analysis of the immune cell compartment in the human small intestine and in Celiac disease. *Nature Communications*, Springer US. 13, 4920. 10.1038/s41467-022-32691-5

[18] Hære, P., Høie, O., Schulz, T., Schönhardt, I., Raki, M. and Lundin, K.E.A. (2016) Long-term mucosal recovery and healing in celiac disease is the rule – not the exception. Scandinavian Journal of Gastroenterology, 51, 1439–46. 10.1080/00365521.2016.1218540

[19] Leonard, M.M., Silvester, J.A., Leffler, D., Fasano, A., Kelly, C.P., Lewis, S.K. et al. (2021) Evaluating Responses to Gluten Challenge: A Randomized, Double-Blind, 2-Dose Gluten Challenge Trial. Gastroenterology, 160, 720-733.e8. 10.1053/j.gastro.2020.10.040

[20] Kutlu, T., Brousse, N., Rambaud, C., Le Deist, F., Schmitz, J. and Cerf-Bensussan, N. (1993) Numbers of T cell receptor (TCR) αβ+ but not of TcR γδ+ intraepithelial lymphocytes correlate with the grade of villous atrophy in coeliac patients on a long term normal diet. Gut, 34, 208–14. 10.1136/gut.34.2.208

[21] Halstensen, T.S., Scott, H. and Brandtzaeg, P. (1989) Intraepithelial T Cells of the TcRγ/δ + CD8 − and Vδ1/Jδ1 + Phenotypes are Increased in Coeliac Disease. Scandinavian Journal of Immunology, 30, 665–72. 10.1111/j.1365-3083.1989.tb02474.x

[22] Mayassi, T., Ladell, K., Gudjonson, H., McLaren, J.E., Shaw, D.G., Tran, M.T. et al. (2019) Chronic Inflammation Permanently Reshapes Tissue-Resident Immunity in Celiac Disease. *Cell*, Elsevier Inc. 176, 967–981.e19. 10.1016/j.cell.2018.12.039

[23] Kurppa, K., Taavela, J., Saavalainen, P., Kaukinen, K. and Lindfors, K. (2016) Novel diagnostic techniques for celiac disease. Expert Review of Gastroenterology & Hepatology, 10, 795–805. 10.1586/17474124.2016.1148599

[24] Taavela, J., Viiri, K., Popp, A., Oittinen, M., Dotsenko, V., Peräaho, M. et al. (2019) Histological, immunohistochemical and mRNA gene expression responses in coeliac disease patients challenged with gluten using PAXgene fixed paraffin-embedded duodenal biopsies. *BMC Gastroenterology*, BMC Gastroenterology. 19, 189. 10.1186/s12876-019-1089-7

[25] Taavela, J., Viiri, K., Välimäki, A., Sarin, J., Salonoja, K., Mäki, M. et al. (2021) Apolipoprotein A4 Defines the Villus-Crypt Border in Duodenal Specimens for Celiac Disease Morphometry. Frontiers in Immunology, 12, 1–10. 10.3389/fimmu.2021.713854

[26] Schumann, M., Siegmund, B., Schulzke, J.D. and Fromm, M. (2017) Celiac Disease: Role of the Epithelial Barrier. *CMGH Cellular and Molecular Gastroenterology and Hepatology*, Elsevier Inc. 3, 150–62. 10.1016/j.jcmgh.2016.12.006

[27] Halstensen, T.S. and Brandtzaeg, P. (1993) Activated T lymphocytes in the celiac lesion: Non-proliferative activation (CD25) of CD4 + α/β cells in the lamina propria but proliferation (Ki-67) of α/β and γ/δ cells in the epithelium. European Journal of Immunology, 23, 505–10. 10.1002/eji.1830230231

[28] Dewar, D.H. and Ciclitira, P.J. (2005) Clinical features and diagnosis of celiac disease. Gastroenterology, 128, 19–24. 10.1053/j.gastro.2005.02.010

[29] Gracz, A.D., Fuller, M.K., Wang, F., Li, L., Stelzner, M., Dunn, J.C.Y. et al. (2013) Brief Report: CD24 and CD44 mark human intestinal epithelial cell populations with characteristics of active and facultative stem cells. Stem Cells, 31, 2024–30. 10.1002/stem.1391

[30] Foroutan, M., Bhuva, D.D., Lyu, R., Horan, K., Cursons, J. and Davis, M.J. (2018) Single sample scoring of molecular phenotypes. *BMC Bioinformatics*, BMC Bioinformatics. 19, 404. 10.1186/s12859-018-2435-4

[31] Abadie, V., Kim, S.M., Lejeune, T., Palanski, B.A., Ernest, J.D., Tastet, O. et al. (2020) IL-15, gluten and HLA-DQ8 drive tissue destruction in coeliac disease. Nature, 578, 600–4. 10.1038/s41586-020-2003-8

[32] Ricaño-Ponce, I., Zhernakova, D. V., Deelen, P., Luo, O., Li, X., Isaacs, A. et al. (2016) Refined mapping of autoimmune disease associated genetic variants with gene expression suggests an important role for non-coding RNAs. Journal of Autoimmunity, 68, 62–74. 10.1016/j.jaut.2016.01.002

[33] Trynka, G., Hunt, K.A., Bockett, N.A., Romanos, J., Mistry, V., Szperl, A. et al. (2011) Dense genotyping identifies and localizes multiple common and rare variant association signals in celiac disease. Nature Genetics, 43, 1193–201. 10.1038/ng.998

[34] Ricaño-Ponce, I., Gutierrez-Achury, J., Costa, A.F., Deelen, P., Kurilshikov, A., Zorro, M.M. et al. (2020) Immunochip meta-analysis in European and Argentinian populations identifies two novel genetic loci associated with celiac disease. European Journal of Human Genetics, 28, 313–23. 10.1038/s41431-019-0520-4

[35] van der Graaf, A., Zorro, M.M., Claringbould, A., Võsa, U., Aguirre-Gamboa, R., Li, C., et al. (2021) Systematic Prioritization of Candidate Genes in Disease Loci Identifies TRAFD1 as a Master Regulator of IFNγ Signaling in Celiac Disease. Frontiers in Genetics, 11, 1–16. 10.3389/fgene.2020.562434

[36] Võsa, U., Claringbould, A., Westra, H.J., Bonder, M.J., Deelen, P., Zeng, B. et al. (2021) Large-scale cis- and trans-eQTL analyses identify thousands of genetic loci and polygenic scores that regulate blood gene expression. Nature Genetics, 53, 1300–10. 10.1038/s41588-021-00913-z

[37] Aguirre-Gamboa, R., de Klein, N., di Tommaso, J., Claringbould, A., Võsa, U., Zorro, M., et al. (2019) Deconvolution of bulk blood eQTL effects into immune cell subpopulations. *BioRxiv*, BMC Bioinformatics. 5, 1–23. 10.1101/548669

[38] Silva-Gomes, R., Mapelli, S.N., Boutet, M., Mattiola, I., Sironi, M., Grizzi, F. et al. (2022) Differential expression and regulation of MS4A family members in myeloid cells in physiological and pathological conditions. Journal of Leukocyte Biology, 111, 817–36. 10.1002/JLB.2A0421-200R

[39] Chen, J., He, R., Sun, W., Gao, R., Peng, Q., Zhu, L. et al. (2020) TAGAP instructs Th17 differentiation by bridging Dectin activation to EPHB2 signaling in innate antifungal response. *Nature Communications*, Springer US. 11. 10.1038/s41467-020-15564-7

[40] Veress, B., Franzén, L., Bodin, L. and Borch, K. (2004) Duodenal intraepithelial lymphocyte-count revisited. Scandinavian Journal of Gastroenterology, 39, 138–44. 10.1080/00365520310007675

[41] Laforenza, U., Miceli, E., Gastaldi, G., Scaffino, M.F., Ventura, U., Fontana, J.M. et al. (2010) Solute transporters and aquaporins are impaired in celiac disease. Biology of the Cell, 102, 457–67. 10.1042/BC20100023

[42] Verbeke, S., Gotteland, M., Fernandez, M., Bremer, J., Rios, G. and Brunser, O. (2002) Basement membrane and connective tissue proteins in intestinal mucosa of patients with coeliac disease. Journal of Clinical Pathology, 55, 440–5. 10.1136/jcp.55.6.440

[43] Leonard, M.M., Bai, Y., Serena, G., Nickerson, K.P., Camhi, S., Sturgeon, C., et al. (2019) RNA sequencing of intestinal mucosa reveals novel pathways functionally linked to celiac disease pathogenesis. Sestak K, editor. PLOS ONE, 14, e0215132. 10.1371/journal.pone.0215132

[44] Koning, F., Schuppan, D., Cerf-Bensussan, N. and Sollid, L.M. (2005) Pathomechanisms in celiac disease. Best Practice and Research: Clinical Gastroenterology, 19, 373–87. 10.1016/j.bpg.2005.02.003

[45] Farstad, I.N., Halstensen, T.S., Lazarovits, A.I., Norstein, J., Fausa, O. and Brandtzaeg, P. (1995) Human Intestinal B-Cell Blasts and Plasma Cells Express the Mucosal Homing Receptor Integrin α4β7. Scandinavian Journal of Immunology, 42, 662–72. 10.1111/j.1365-3083.1995.tb03709.x

[46] Yu, X., Vargas, J., Green, P.H.R. and Bhagat, G. (2021) Innate Lymphoid Cells and Celiac Disease: Current Perspective. *Cmgh*, Elsevier Inc. 11, 803–14. 10.1016/j.jcmgh.2020.12.002

[47] Bragde, H., Jansson, U., Fredrikson, M., Grodzinsky, E. and Söderman, J. (2018) Celiac disease biomarkers identified by transcriptome analysis of small intestinal biopsies. Cellular and Molecular Life Sciences, Springer International Publishing. 75, 4385–401. 10.1007/s00018-018-2898-5

[48] Yasuda, K., Nakanishi, K. and Tsutsui, H. (2019) Interleukin-18 in Health and Disease. International Journal of Molecular Sciences, 20, 649. 10.3390/ijms20030649

[49] León, A.J., Garrote, J.A., Blanco-Quirós, A., Calvo, C., Fernández-Salazar, L., Del Villar, A. et al. (2006) Interleukin 18 maintains a long-standing inflammation in coeliac disease patients. Clinical and Experimental Immunology, 146, 479–85. 10.1111/j.1365-2249.2006.03239.x

[50] Nowarski, R., Jackson, R., Gagliani, N., de Zoete, M.R., Palm, N.W., Bailis, W., et al. (2015) Epithelial IL-18 Equilibrium Controls Barrier Function in Colitis. Cell, 163, 1444–56. 10.1016/j.cell.2015.10.072

[51] Jansen, R., Hottenga, J.J., Nivard, M.G., Abdellaoui, A., Laport, B., de Geus, E.J., et al. (2017) Conditional eQTL analysis reveals allelic heterogeneity of gene expression. Human Molecular Genetics, 26, 1444–51. 10.1093/hmg/ddx043

[52] Das, S., Forer, L., Schönherr, S., Sidore, C., Locke, A.E., Kwong, A. et al. (2016) Next-generation genotype imputation service and methods. Nature Genetics, 48, 1284–7. 10.1038/ng.3656

[53] Kim, D., Langmead, B. and Salzberg, S.L. (2015) HISAT: a fast spliced aligner with low memory requirements. Nature Methods, 12, 357–60. 10.1038/nmeth.3317

[54] Li, H., Handsaker, B., Wysoker, A., Fennell, T., Ruan, J., Homer, N. et al. (2009) The Sequence Alignment/Map format and SAMtools. Bioinformatics, 25, 2078–9. 10.1093/bioinformatics/btp352

[55] Anders, S., Pyl, P.T. and Huber, W. (2015) HTSeq-A Python framework to work with high-throughput sequencing data. Bioinformatics, 31, 166–9. 10.1093/bioinformatics/btu638

[56] Andrews, S. (2010) FastQC a Quality Control Tool for High Throughput Sequence Data [Internet].

[57] (2019) Picard Toolkit [Internet]. Broad Institute.

[58] Love, A.M., Anders, S., Huber, W. and Love, M.M. (2017) Package ‘ DESeq2.’

[59] Gu, Z. (2022) Complex heatmap visualization. IMeta, 1. 10.1002/imt2.43

[60] Mächler, M., Rousseeuw, P., Struyf, A., Hubert, M. and Hornik, K. (2012) Cluster: Cluster Analysis Basics and Extensions. R Packag.

[61] Kassambara, A. and Mundt, F. (2020) factoextra: Extract and Visualize the Results of Multivariate Data Analyses [Internet].

[62] Fabregat, A., Jupe, S., Matthews, L., Sidiropoulos, K., Gillespie, M., Garapati, P., et al. (2018) The Reactome Pathway Knowledgebase. Nucleic Acids Research, Oxford University Press. 46, D649–55. 10.1093/nar/gkx1132

[63] Yu, G., Wang, L.G., Han, Y. and He, Q.Y. (2012) ClusterProfiler: An R package for comparing biological themes among gene clusters. OMICS A Journal of Integrative Biology, 16, 284–7. 10.1089/omi.2011.0118

[64] Robinson, M.D., McCarthy, D.J. and Smyth, G.K. (2010) edgeR: a Bioconductor package for differential expression analysis of digital gene expression data. Bioinformatics, 26, 139–40. 10.1093/bioinformatics/btp616

[65] Zhernakova, D. V, Deelen, P., Vermaat, M., van Iterson, M., van Galen, M., Arindrarto, W., et al. (2017) Identification of context-dependent expression quantitative trait loci in whole blood. Nature Genetics, 49, 139–45. 10.1038/ng.3737

[66] R Core Team. (2019) R: A Language and Environment for Statistical Computing [Internet]. Vienna, Austria.

[67] Wickham, H. (2016) ggplot2: Elegant Graphics for Data Analysis [Internet]. Springer-Verlag New York.

