## Supplementary Figures for "Gene expression and eQTL analysis reflect the heterogeneity in the inflammatory status of the duodenal epithelial lining in coeliac disease"

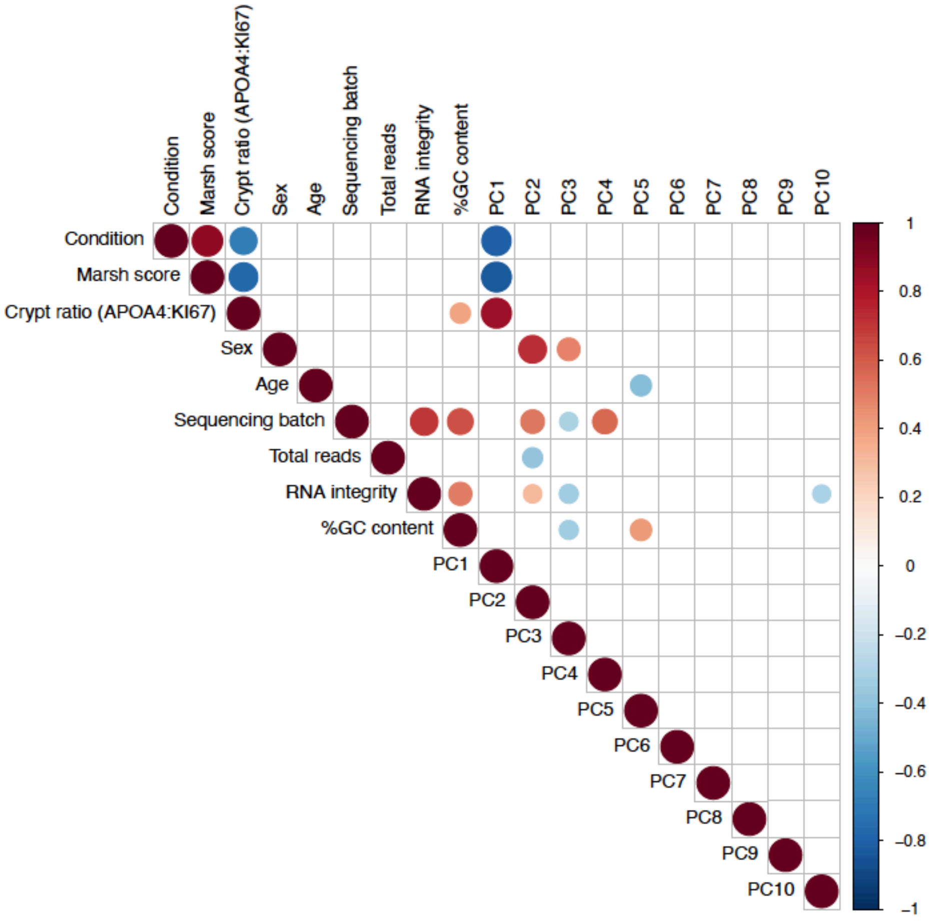


**Supplementary Figure 1.** Spearman correlations between variables and PCs in dataset (p-value < 0.01).

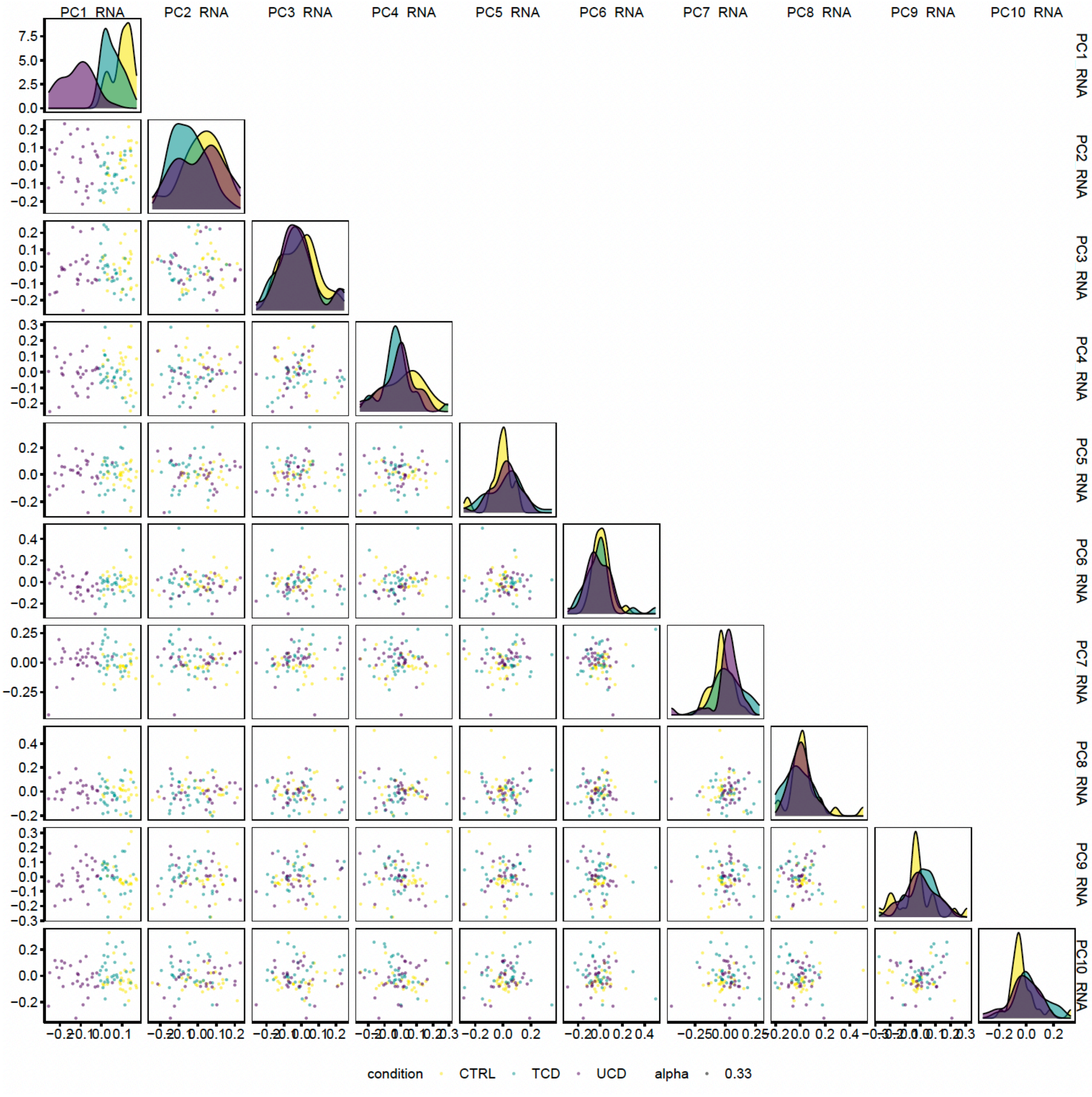


**Supplementary Figure 2.** Comparison of PCs in relation to CeD condition.

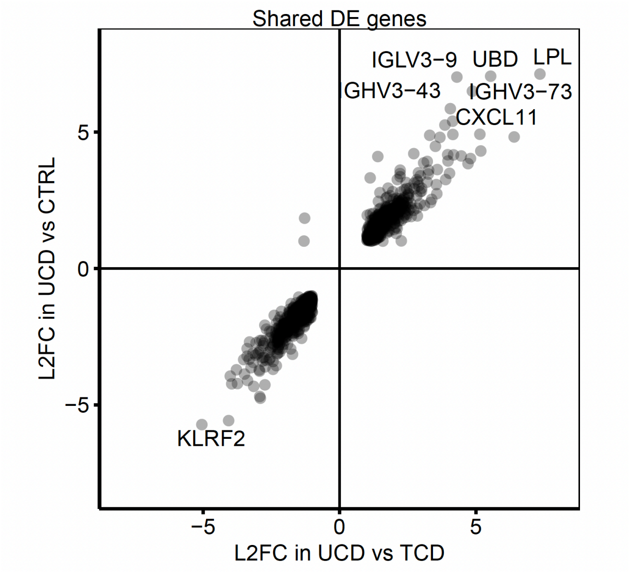


**Supplementary Figure 3.** Concordance between the DE gene results of UCD vs CTRL and UCD vs TCD.

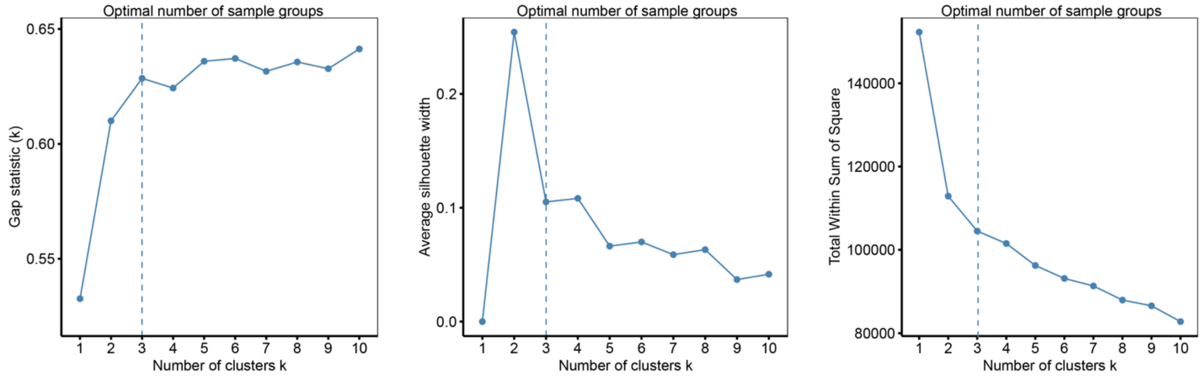


**Supplementary Figure 4.** Results of the k-Means clustering of samples. Exploration of the optimal number of clusters of samples. The cluster number (k) of sample groups was determined by comparing the results of three different methods: gap statistic (500 permutations), average silhouette width, and total within sum-of-square when clustering from 1 to 10 groups. Dotted line indicates the optimal number of cluster of samples determined for each test.

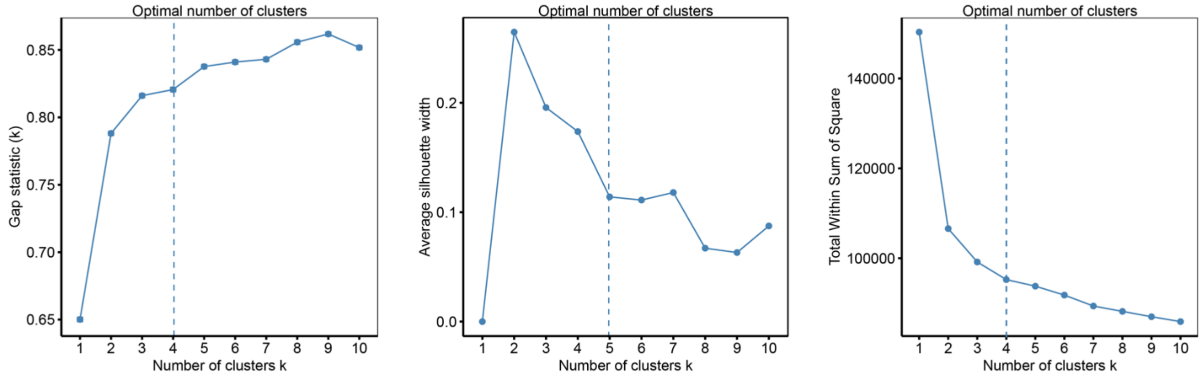


**Supplementary Figure 5.** Results of k-Means clustering analysis of DE genes. Exploration of optimal number of clusters of DE genes. Cluster number (k) of DE genes was determined by comparing the results of three different methods: gap statistic (500 permutations), average silhouette width, and total within sum-of-square when clustering from 1 to 10 groups. Dotted line indicates the optimal number of clusters determined for each test.

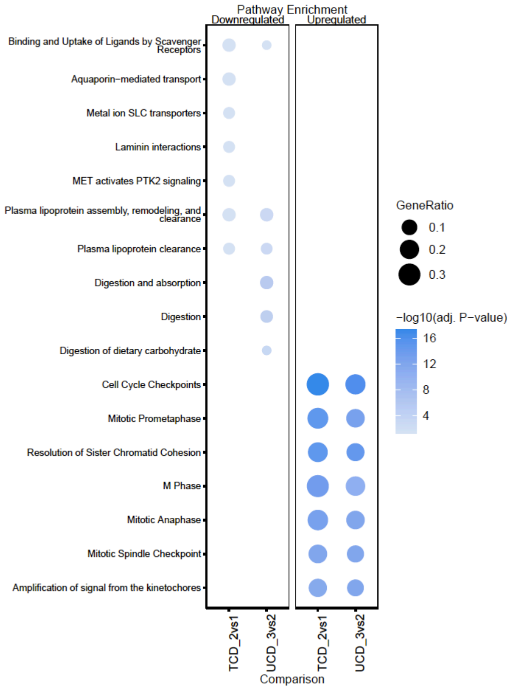


**Supplementary Figure 4.** Enrichment analysis of inter-variation within TCD and UCD.

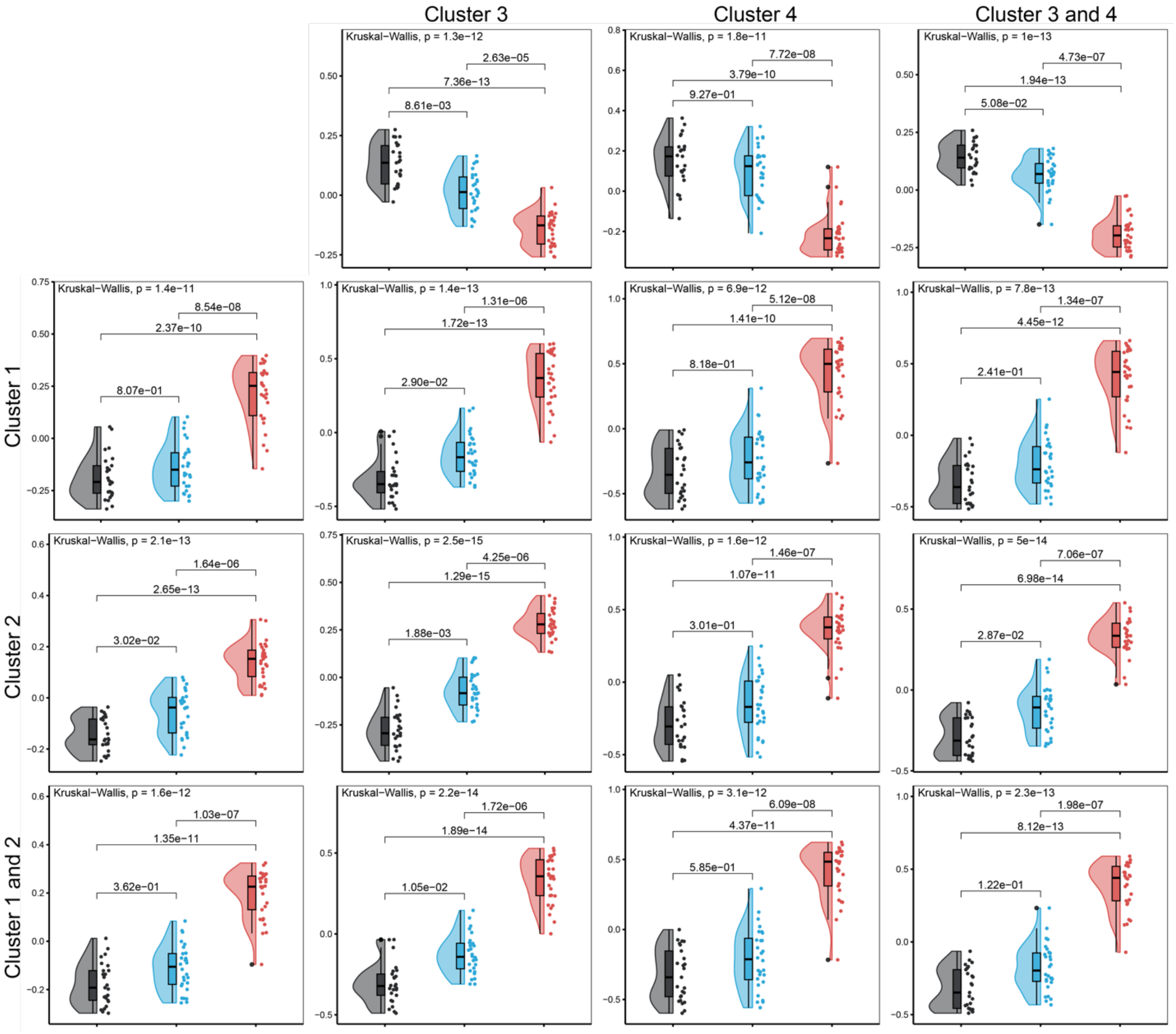


**Supplementary Figure 5.** Singscores of samples using clusters of DE genes. Singscores (y-axis) of samples obtained using genes of cluster 3, cluster 4, or both combined as an upregulated set (top) and/or genes from cluster 1, cluster 2, or both combined as a downregulated set (left). Samples are divided per disease condition: CTRL (grey), TCD (blue), and UCD (red). Adjusted p-value < 0.01, Dunn test, Bonferroni correction.

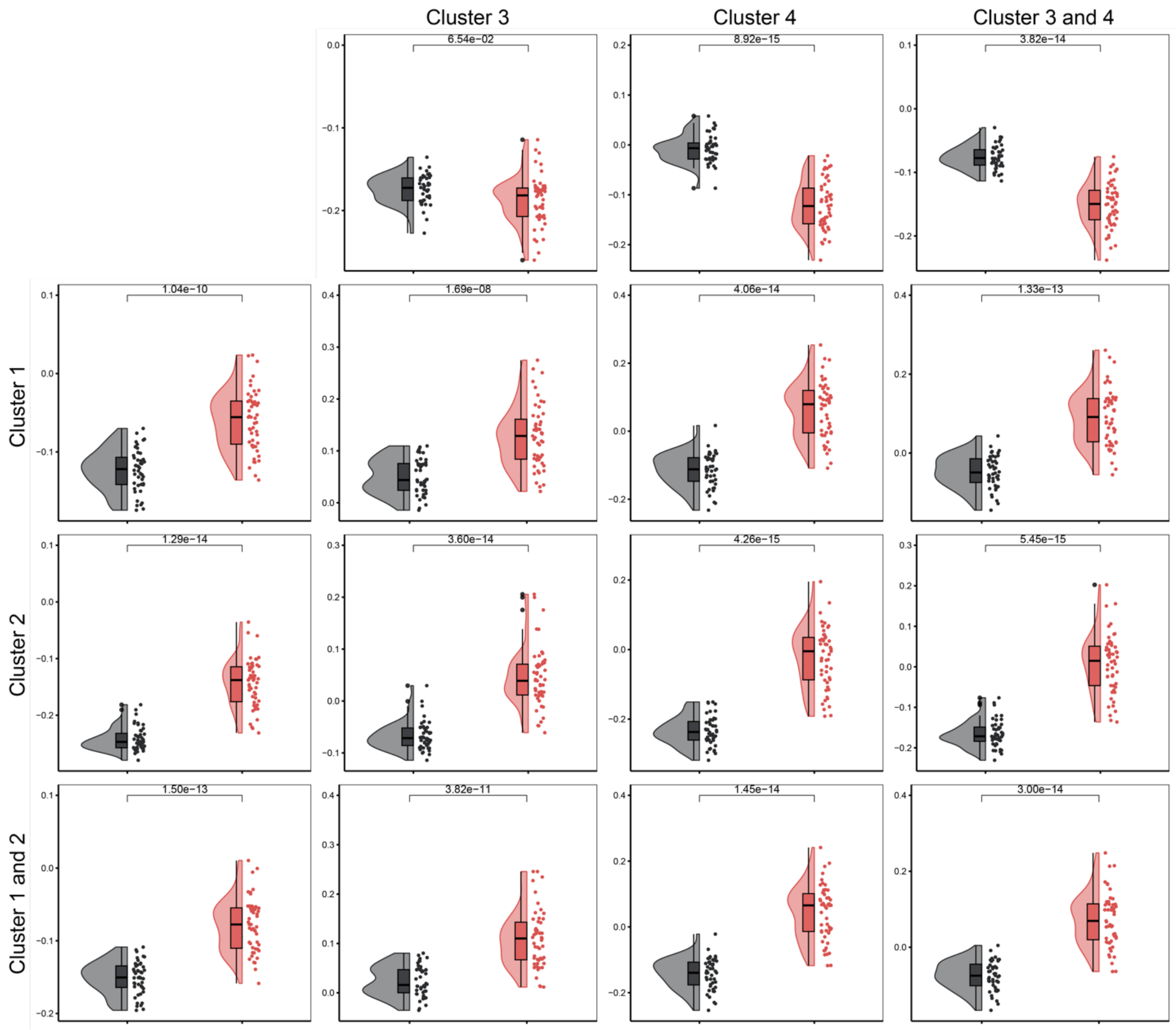


**Supplementary Figure 6.** Singscores of an external cohort using clusters of DE genes. Singscores (y-axis) of the external cohort obtained by using genes of cluster 3, cluster 4, or both combined as an upregulated set (top) and/or genes from cluster 1, cluster 2, or both combined as a downregulated set (left). Samples are divided per disease condition: CTRL (grey) and UCD (red). Adjusted p-value < 0.01, Dunn test, Bonferroni correction.
